# Coordinated demethylation of H3K9 and H3K27 is required for rapid inflammatory responses of endothelial cells

**DOI:** 10.1101/456491

**Authors:** Yoshiki Higashijima, Yusuke Matsui, Teppei Shimamura, Shuichi Tsutsumi, Ryo Nakaki, Yohei Abe, Verena M. Link, Mizuko Osaka, Masayuki Yoshida, Ryo Watanabe, Toshihiro Tanaka, Akashi Taguchi, Mai Miura, Tsuyoshi Inoue, Masaomi Nangaku, Hiroshi Kimura, Tetsushi Furukawa, Hiroyuki Aburatani, Youichiro Wada, Christopher K. Glass, Yasuharu Kanki

**Affiliations:** Department of Bioinformational Pharmacology, Tokyo Medical and Dental University, Tokyo 113-8510, Japan; Isotope Science Center, The University of Tokyo, Tokyo 113-0032, Japan; Department of Systems Biology, Graduate School of Medicine, Nagoya University, Nagoya 466-8550, Japan; Division of Genome Sciences, RCAST, The University of Tokyo, Tokyo 153-8904, Japan; Department of Cellular and Molecular Medicine, School of Medicine, University of California, San Diego, La Jolla, CA 92093, USA; Faculty of Biology, Division of Evolutionary Biology, Ludwig-Maximilian University of Munich, Munich, Germany; Department of Nutrition in Cardiovascular Disease, Tokyo Medical and Dental University, Yushima, Bunkyo-ku, Tokyo 113-8510, Japan; Department of Life Sciences and Bioethics, Tokyo Medical and Dental University, Yushima, Bunkyo-ku, Tokyo 113-8510, Japan; Department of Human Genetics and Disease Diversity, Graduate School of Medical and Dental Sciences, Tokyo Medical and Dental University, 113-8510 Tokyo, Japan; Bioresource Research Center, Tokyo Medical and Dental University, 113-8510 Tokyo, Japan; Division of Nephrology and Endocrinology, The University of Tokyo Graduate School of Medicine, 113-0033 Tokyo, Japan; Cell Biology Center, Institute of Innovative Research, Tokyo Institute of Technology, 226-8503 Yokohama, Japan; Department of Medicine, University of California, San Diego, La Jolla 92093, CA, USA

**Keywords:** Inflammation, Atherosclerosis, histone modification enzyme, repressive histone mark, super enhancer, chromatin architecture, chromatin conformation change

## Abstract

Lysine 9 di-methylation and lysine 27 tri-methylation of histone H3 (H3K9me2 and H3K27me3) are generally linked to gene repression. However, the functions of repressive histone methylation dynamics during inflammatory responses remain enigmatic. We found that tumor necrosis factor (TNF)-α rapidly induces the co-occupancy of lysine demethylases 7A (KDM7A) and 6A (UTX) with nuclear factor kappa-B (NF-κB) recruited elements in human endothelial cells. KDM7A and UTX demethylate H3K9me2 and H3K27me3, respectively, and both are required for activation of NF-κB-dependent inflammatory genes. Chromosome conformation capture-based methods demonstrated increased interactions between TNF-α-induced super enhancers at NF-κB-relevant loci, coinciding with KDM7A- and UTX-recruitment. Simultaneous inhibition of KDM7A and UTX significantly reduced leukocyte adhesion in mice, establishing the biological and potential translational relevance of this mechanism. Collectively, these findings suggest that rapid erasure of repressive histone marks by KDM7A and UTX is essential for NF-κB-dependent regulation of genes that control inflammatory responses of endothelial cells.

**HIGHLIGHTS:**

1. KDM7A and UTX cooperatively control NF-κB-dependent transcription in vascular endothelial cells.
2. Demethylation of repressive histone marks by KDM7A and UTX is critical for early inflammatory responses.
3. KDM7A and UTX are associated with TNF-α-induced looping of super enhancers.
4. Pharmacological inhibition of KDM7A and UTX reduces leukocyte adhesive interactions with endothelial cells in mice.

## INTRODUCTION

Precise control of inflammation is essential for the immune system and the maintenance of tissue homeostasis in higher eukaryotes. Insufficient inflammatory responses increase the risk of contracting infectious diseases, while excessive or inadequate responses contribute to many common and life-threatening illnesses. Atherosclerosis is an example of a chronic low-grade inflammatory disease of arteries, mediated by the accumulation of plaque within the blood vessel walls. Heart attack and stroke as a sequel of atherosclerosis are responsible for an immense burden of morbidity and mortality (Benjamin et al., 2017). Endothelial cells (ECs) are not only static cells that are components of the blood vessel system, but also play pivotal roles in the development of atherosclerosis. Pro-inflammatory stimuli, such as hemodynamic turbulence, hyperlipidemia, and inflammatory cytokines, induce the expression of adhesion molecules on the luminal EC surface, which causes leukocyte recruitment and a subsequent cascade of inflammatory processes (Gistera and Hansson, 2017; Libby, 2002; Libby et al., 2011). Although the aberrant expression of adhesion molecules has been implicated in the initiation step of atherosclerosis, the molecular mechanisms responsible for the induction of adhesion molecules are currently incompletely understood.

Recent technological advances in next generation sequencing (NGS) have revealed that epigenetic modifications, including DNA methylation, histone modification, non-coding RNA, and chromosomal conformation, influence gene expression, thus contributing to the regulation of cell fate, the maintenance of cell specificity, and cell type-specific functions (Roadmap Epigenomics et al., 2015). In vascular development, histone modifications correlate with master transcription factors during EC differentiation from embryonic stem (ES) cells (Kanki et al., 2017). In addition, to maintain EC specificity, the transcription factor GATA2 regulates EC-specific gene expression, correlated with the cell type-specific chromatin conformation and histone modifications (Kanki et al., 2011). In ECs, several miRNAs reportedly regulate the balance of pro- or anti-inflammatory signaling pathways in response to external stimuli, including oscillatory flow and inflammatory cytokines (Feinberg and Moore, 2016). Consistently, we previously reported that tumor necrosis factor (TNF)-α-responsive miRNAs not only co-associate with TNF-α-responsive coding genes but also with genes harboring miRNAs, suggesting that transcriptional factories specialized to produce non-coding transcripts control the inflammatory responses (Papantonis et al., 2012). Our group has also reported that interleukin-4 induced adhesion molecules, through transcription factor STAT6-mediated histone modification changes, in cultured human ECs (Tozawa et al., 2011). Taken together, epigenetic regulations in ECs play critical roles in the initiation and progression of vascular related diseases, including atherosclerosis.

The post-transcriptional modifications (methylation, acetylation, and ubiquitination) of histone tails correlate with the chromatin state of either activated or repressed transcription of the associated genes. In particular, di-methylation of histone H3 at lysine 9 (H3K9me2) and tri-methylation of histone H3 at lysine 27 (H3K27me3) are mostly associated with gene repression. Several *in vitro* differentiation studies have demonstrated that the jumonji C (JmjC) domain containing protein, ubiquitously transcribed tetratricopeptide repeat, X linked (UTX), also known as KDM6A, demethylates H3K27me3 and is required for the activation of specific gene expression during lineage commitment (Agger et al., 2007; Lan et al., 2007). In addition, exome- and genome-wide sequencing strategies have identified UTX mutations and deletions in multiple cancer types, including leukemia (Bailey et al., 2016; Cancer Genome Atlas Research et al., 2013; Dalgliesh et al., 2010; Gui et al., 2011; Huether et al., 2014; Ntziachristos et al., 2014; van Haaften et al., 2009), indicating the possible role of UTX as a tumor suppressor in cancer biology. Another JmjC domain containing protein, lysine demethylase 7A (KDM7A), also known as JHDM1D, contains a plant homeo domain (PHD) and demethylates H3K9me2 and H3K27me2. Furthermore, some groups have reported that KDMA7A functions as an eraser of silencing marks on chromatin, mostly during brain development, cell differentiation, cell cycle, and cell proliferation (Horton et al., 2010; Osawa et al., 2011; Tsukada et al., 2010). Although the significance of histone demethylases in transcriptional control during development and in cancer has been extensively studied, fewer studies have examined the potential roles of histone demethylases in acute responses, including inflammation in terminally differentiated cells (De Santa et al., 2009; De Santa et al., 2007; Kruidenier et al., 2012).

Inflammatory responses are controlled by master transcription factors, including Nuclear factor-kappa B (NF-κB) (Zhang et al., 2017). NF-κB cooperatively interacts at *cis* regulatory elements (*i.e*., proximal promoters and distal enhancers) in a context-dependent manner to elicit the appropriate gene expression profiles. NF-κB binding is coupled with the dynamic remodeling of the epigenetic landscape and chromatin structure, to change global gene expression patterns responsible for cell fate, phenotype, and behavior. Previous studies of transcription factor binding profiles in different cell types and conditions revealed that most binding occurs in intronic or intergenic regions, showing that enhancers play crucial roles in determining the cell type-specific transcriptome (Adam et al., 2015; Heinz et al., 2015; Saint-Andre et al., 2016). Other studies demonstrated the importance of super enhancers (SEs), also known as stretch or large enhancer clusters, for the regulation of genes correlated with cell identity (Hnisz et al., 2013; Parker et al., 2013; Suzuki et al., 2017; Whyte et al., 2013). The expression of genes associated with SEs is particularly sensitive to outside stimuli, including inflammatory cytokines, which may control and facilitate cell state transitions (Brown et al., 2014; Hah et al., 2015; Schmidt et al., 2015). In human ECs, TNF-α-induced SEs display extremely high occupancies of NF-κB, active histone marks (H3K27ac), and the epigenetic reader protein bromodomain-containing protein 4 (BRD4). The pharmacological inhibition of BRD4 by JQ1 inhibited a broad range of TNF-α-responsive genes, suggesting the important role of SEs in the inflammatory response of ECs (Brown et al., 2014). More recently, Beyaz et al. showed that UTX facilitates the accessibility of SEs that establish the cell identity of invariant natural killer T cells, in accordance with low levels of H3K27me3 (Beyaz et al., 2017). This study raised the possibility that the erasure of repressive histone marks by histone demethylases correlates with the formation of SEs and the activation of associated genes; however, these points have not been addressed during the early inflammatory responses.

Recent advances in chromosome conformation capture (3C)-based technologies have emphasized the relevance of the three-dimensional (3D) genome organization for transcriptional control. Chromatin interaction analysis with paired-end tag sequencing (ChIA-PET) (Fullwood et al., 2009) and HiChIP (Mumbach et al., 2016) have identified chromatin loops (*e.g*., promoter-enhancer interactions) mediated by specific protein factors, which are functionally associated with gene expression programs. Hi-C (Lieberman-Aiden et al., 2009) has also revealed that the genome is organized into topology associating domains (TADs) of several hundred kilobases to a few megabases, which play important roles in genome organization and proper control of gene expression. Megabase TADs are reportedly already formed in ES cells and largely conserved through development, among different cell types and in response to specific stimuli (Dixon et al., 2012; Jin et al., 2013), although recent studies have suggested that submegabase (sub)-TADs could have plastic properties (Kim et al., 2018; Ogiyama et al., 2018; Phillips-Cremins et al., 2013; Siersbaek et al., 2017). Furthermore, using 3C, circular 3C (4C), and ChIA-PET, we showed that human ECs have cell type-specific small chromatin loops, which correlate with gene expression profiles (Inoue et al., 2014; Kanki et al., 2011; Mimura et al., 2012; Papantonis et al., 2012). However, there is no clear consensus on the plasticity of chromatin interactions (*i.e*., chromatin loops or small TADs) in response to external stimuli, particularly in terminally differentiated cells.

In this study, we identified a novel TNF-α-responsive miRNA, miR-3679-5p, which inhibits the induction of adhesion molecules by targeting two histone demethylases, KDM7A and UTX, in cultured human ECs. Importantly, an RNA-seq analysis revealed that KDM7A and UTX cooperatively control the expression of not only adhesion molecules but also many other NF-κB-dependent genes. Consistent with this, TNF-α stimuli recruited KDM7A and UTX to the NF-κB-related elements, where the repressive histone marks H3K9me2 and H3K27me3 were rapidly removed. Interestingly, TNF-α-induced SE-regions displayed high occupancies of KDM7A and UTX, which were enriched for cardiovascular disease-associated genetic variants. Furthermore, Hi-C in combination with ChIA-PET revealed that TNF-α-responsive SE-SE interactions were newly formed within sub-TADs, immediately following TNF-α stimulation. Finally, the pharmacological inhibition of KDM7A and UTX led to the reduction of leukocyte adhesive interactions on the surfaces of TNF-α-activated ECs in mice. Taken together, these data suggest that erasing repressive histone marks by histone demethylases within NF-κB-related regions could be a cue during inflammatory responses in ECs. Our findings about the synergistic functions of these two histone demethylases give new insights into the initiation of atherosclerosis and may provide therapeutic opportunities for the treatments of vascular inflammatory diseases.

## RESULTS

### A novel miRNA, miR-3679-5p, inhibits monocyte adhesion to ECs

To explore the transcriptional regulatory dynamics that occur during the inflammatory activation of ECs, we activated human umbilical vein ECs with 10 ng/ml of TNF-α, a canonical pro-inflammatory stimulus, for 4 and 24 hrs and analyzed the whole transcriptome by NGS. As a result, the TNF-α-upregulated genes (656; more than 2-fold by TNF-α treatment) can be divided into two classes, those that are strongly induced at 4 h (class 1) and those that are strongly induced at 24 h (class 2). Consistent with our previous report (Tozawa et al., 2011), important adhesion molecules involved in monocyte recruitment to atherosclerotic lesions, including vascular cell adhesion molecule 1 (VCAM1), intracellular adhesion molecule 1 (ICAM1), and E-selectin (SELE), were classified into class 1 (Figure 1A, Figure S1A-D, and Table S1). A gene ontology (GO) analysis further revealed that the class 1 and class 2 genes were commonly related to inflammatory responses, whereas the biological processes of extracellular matrix regulation were highly enriched in class 2 (Figure S1E-F). Interestingly, TNF-α-downregulated genes (501; less than 0.5-fold by TNF-α treatment) were associated with developmental pathways, including angiogenesis and blood vessel morphogenesis (Figure S1G and H, and Table S1). These data led us to speculate that during the rapid activation of terminally differentiated ECs, the inflammatory stimulation up-regulated the transcription of genes necessary for the cytokine response and repressed the genes important for the development of ECs, possibly because of the limitations of ATP and active RNA polymerase II (RNAP).

**Figure 1.**
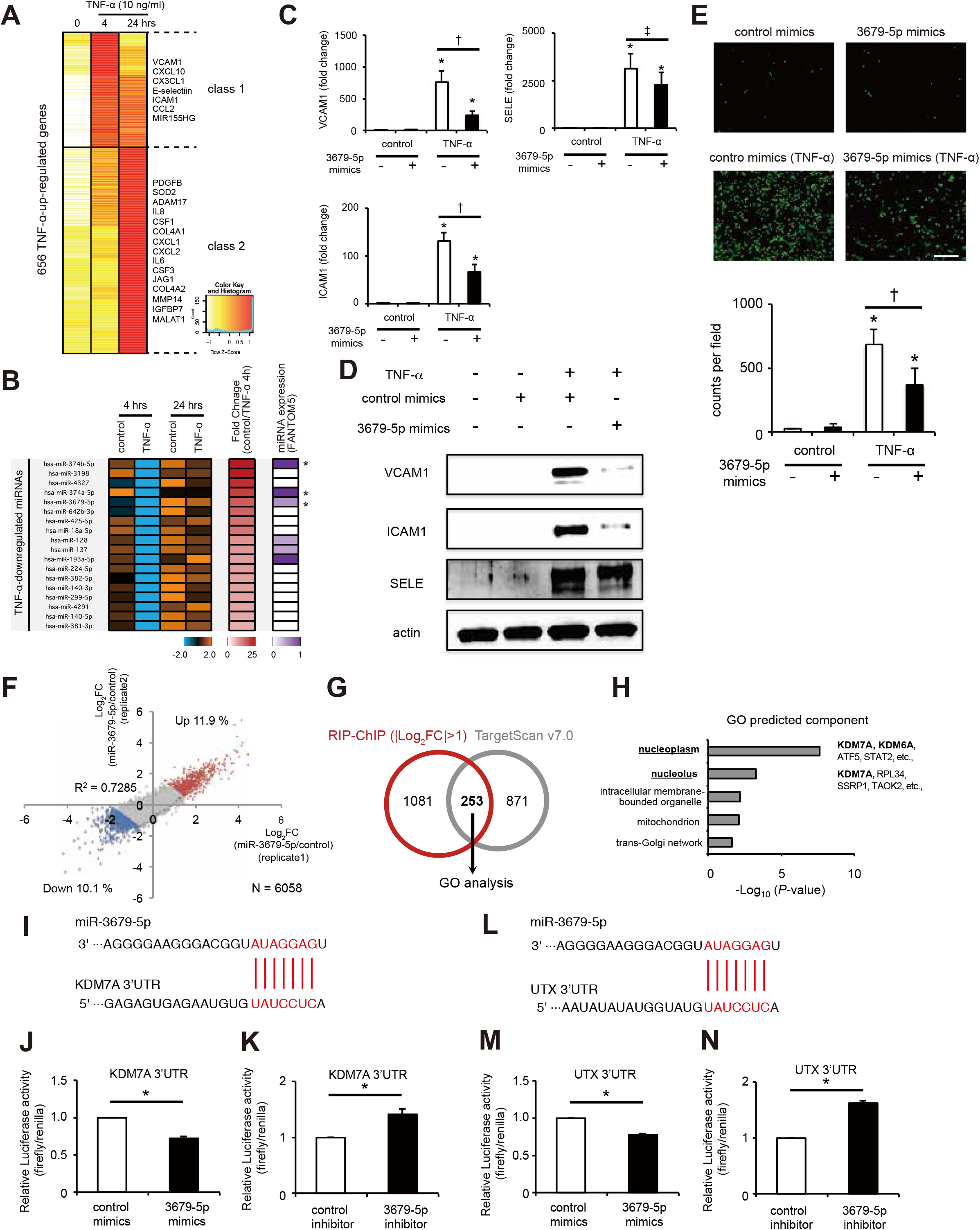
The novel miR-3679-5p suppresses monocyte adhesion and targets two histone demethylases, KDM7A and UTX, in human inflammatory activated ECs. (A) Heat map representation of the TNF-α-up-regulated genes. Genes were classified based on the TNF-α-treated time points, using the algorithm “HOPACH”. (B) Heat map showing down-regulated miRNAs at 4 hrs, but not 24 hrs, after TNF-α treatment. The middle column indicates the fold-change values (control/TNF) at 4 hrs. The right column indicates the relative expression level of each miRNA from the FANTOM5 database (FANTOM Consortium et al., 2014). (C) Bar plots showing mean mRNA levels of vascular cell adhesion molecule-1 (VCAM1), intracellular adhesion molecule-1 (ICAM1), and E-selectin (SELE), measured 4 hrs after stimulation of human endothelial cells (ECs) with or without TNF-α ± miR-3679-5p. Data are shown as means ± SE. ^*^*P* < 0.05 compared to TNF-α (-) mimics (-). † *P* < 0.05 and ‡ *P* = 0.203 compared to TNF-α (+) mimics (-). (D) Western blots for VCAM1, ICAM, SELE, and β-actin in lysates from ECs treated with or without TNF-α (4 hrs) ± miR-3679-5p. (E) Representative images (top) and bar plot quantification (bottom) showing the adhesion of calcein-labeled U937 monocytes to ECs treated with or without TNF-α ± miR-3679-5p. Scale bar represents 250 μm. Data are shown as means ± SD. ^*^*P* < 0.05 compared to TNF-α (-) mimics (-). † *P* < 0.05 compared to TNF-α (+) mimics (-). (F) The scatter plot shows the correlation of RNA immunoprecipitation (RIP) followed by microarray data from two replicate screens for the identification of miR-3679-5p target genes. Genes up- and down-regulated by miR-3679-5p treatment (|average Log_2_FC|>1) are indicated in red and blue, respectively. (G) Venn diagram showing the overlap of the genes selected by RIP-ChIP (1,334) and TargetScan7.0 (1,224) (http://www.targetscan.org/vert_70/). (H) Gene ontology (GO) analysis of the 253 potential target genes for miR-3679-5p, as determined in Figure 1G. The *P*-values of each category analyzed from DAVID (https://david.ncifcrf.gov/) are shown in the bar graphs. (I) Seed sequence of miR-3679-5p and complementary 3′ UTR sequence of lysine demethylase 7A (KDM7A). The letters in blue indicate matched bases. (J and K) Effect of miR-3679-5p mimics (J) or miR-3679-5p inhibitor (K) on luciferase activity in pmirGLO-transfected ECs expressing the 3′ UTR of KDM7A. Data are shown as means ± SE. ^*^*P* < 0.05 compared to control. (L) Seed sequence of miR-3679-5p and complementary 3′ UTR sequence of lysine demethylase 6A (UTX). The letters in blue indicate matched bases. (M and N) The effect of miR-3679-5p mimics (M) or miR-3679-5p inhibitor (N) on luciferase activity in pmirGLO-transfected ECs expressing the 3′ UTR of UTX was measured. Data are shown as means ± SE. ^*^*P* < 0.05 compared to control.

In our previous study, we noticed that the downregulation of miRNAs is coupled with the upregulation of their corresponding mRNAs and vice versa after inflammatory stimuli. (Papantonis et al., 2012). Therefore, we hypothesized that the miRNAs intermediate rapid response of adhesion molecule mRNAs would increase, and performed miRNA-microarray analyses of ECs at baseline and treated with TNF-α for 4 and 24 hrs. Probes with low expression values (raw signal < 0.1 in all conditions) were removed, and 199 miRNAs were identified in total (Table S1). Based on the hypothesis that the expression levels of miRNAs that regulate adhesion molecules in class 1 of Figure 1A change after 4 hrs of TNF-α treatment, we selected TNF-α-responsive miRNAs (> 2-fold expression change at 4 hrs relative to control). Interestingly, many miRNAs were down-regulated in response to TNF-α treatment (Figure 1B) although TNF-α increased some miRNAs, including miR-155 and miR-31, which were previously reported to be up-regulated by TNF-α treatment (Table S1) (Papantonis et al., 2012; Suarez et al., 2010). Of 31 TNF-α-down-regulated miRNAs at 4 hrs, 18 miRNAs were unchanged at 24 hrs with or without TNF-α treatment (Figure 1B). Among them, 6 miRNAs are expressed in human ECs according to the FANTOM5 database, a collaborative omics data integration and interactive visualization system (de Rie et al., 2017). We further selected the top three most down-regulated miRNAs (miR-3679-5p, miR-374b-5p, miR374a-5p) and examined the effects of these miRNAs on the TNF-α-induced mRNA expression of adhesion molecules. Treatment of the cells with miR-3679-5p only significantly decreased TNF-α-induced mRNA expression of VCAM1 (Figure 1C, and Figure S1I and J). Interestingly, miR-3679-5p is human-specific and not conserved in mammals, and it reduced not only the mRNA induction of VCAM1, but also those of ICAM1 and SELE (Figure 1C and 1D). Moreover, to test whether the reduction of these three adhesion molecules by miR-3679-5p overexpression influenced cell adhesion, we performed a monocyte adhesion assay. As shown in Figure 1E, the overexpression of miR-3679-5p led to a reduction of monocyte binding to cultured ECs under TNF-α stimulation. Taken together, these results suggested that miR-3679-5p could be a miRNA that potentially regulates adhesion molecules during TNF-α-induced early inflammatory responses in ECs.

### Identification of KDM7A and UTX as potential target genes of miR-3679-5p

MiRNAs function as gene repressors by complementary base pairing with their target mRNAs within miRNA-mRNA duplexes. Each miRNA has many potential target mRNAs, because its ‘seed’ sequence contains ~6 to ~8-nt (Garcia et al., 2011). To identify the target mRNAs of miR-3679-5p in human ECs, we performed biologically duplicate RNA immunoprecipitation (RIP)-Chip analyses using an antibody against argonaute 2 (AGO2), an essential component of the RNA-induced silencing complex (RISC). The miRNA that binds to its target mRNA is incorporated into a RISC (Bartel, 2018), and can therefore be co-isolated with an antibody against AGO2. Utilizing this approach, ECs transfected with miR-3679-5p or negative control miRNA mimics were stimulated with TNF-α and subsequently immunoprecipitated with an anti-AGO2 antibody. The AGO2-bound RNAs were subjected to a microarray analysis and the differentially regulated mRNAs were filtered out (>2-fold different between miR-3679-5p and negative control in both replicates), identifying a total of 1,334 probes (Figure 1F and G, Table S1). To detect the direct target mRNAs, the 1,334 probes were further filtered by the computational algorithm, TargetScan7.0 (Agarwal et al., 2015), and 253 genes were identified as bona fide targets for miR-3679-5p (Figure 1G and Table S1). To fully inspect the functions of these 253 genes, we performed a GO analysis. The most significantly enriched GO term was nucleoplasm, consisting of 66 genes (Figure 1H, *P* = 6.22E-06 after Benjamini correction; Table S1). Interestingly, these 66 genes included the two histone demethylases KDM7A and KDM6A (UTX), which modify the methylation levels of histone tails and participate in gene regulation (Agger et al., 2007; Lan et al., 2007; Tsukada et al., 2010). Since we previously reported that histone modification changes of H3K4me1 and H3K4me3 preceded the mRNA induction of genes associated with inflammatory responses in ECs (Tozawa et al., 2011), we focused on these two histone modification enzymes in this work. Using qRT-PCR, we found that the expression of KDM7A and UTX in ECs transfected with miR-3679-5p was significantly downregulated, in comparison to cells transfected with negative control miRNA mimics (data not shown; *P* = 0.0001 and 0.0351, respectively). Consistently, the 3′UTRs of KDM7A and UTX have seed sequences corresponding to miR-3679-5p (Figure 1I and L). Furthermore, a reporter assay confirmed that the transfection of mimics and the inhibition of miR-3679-5p suppressed and increased the luciferase activities, respectively, in ECs (Figure 1J, K, M and N), suggesting that KDM7A and UTX are potentially the direct targets of miR-3679-5p.

### KDM7A and UTX participate in TNF-α-induced NF-κB-p65 signaling pathways leading to activation of ECs

To examine whether KDM7A and UTX are involved in the induction of TNF-α-responsive genes and the activation of the NF-κB pathway, we performed an RNA-seq analysis in KDM7A- and UTX-knockdown ECs at 4 hrs after the TNF-α treatment (Figure S2A-D). The TNF-α-responsive genes were identified as those that were up-regulated by more than 2-fold after TNF-α treatment. We identified a total of 404 genes as TNF-α-responsive genes, of which 123 genes were down-regulated after NF-κB inhibition by IκK-α (BAY 11-7082, less than 0.75-fold vs. TNF-α, Table S2). Since NF-κB is a master regulator of gene transcription during inflammatory processes and plays a critical role in EC activation (Brown et al., 2014; Papantonis et al., 2012), the 123 NF-κB-dependent genes were further analyzed by clustering their siRNA-treated expression profiles and divided into five clusters (Figure 2A). The siRNA-mediated knockdown of KDM7A down-regulated almost half of the NF-κB-dependent genes (*e.g*., VCAM1, CCL2, CXCL2, CXCL3, CX3CL1, IL8, and SOD2), which play important roles in the immune response and cell chemotaxis (class 2, Figure 2A). The siRNA knockdown of UTX down-regulated a small number of NF-κB-dependent genes, including SELE (class 3, Figure 2A). Several other NF-κB-dependent genes were down-regulated by both siKDM7A and siUTX (class 1, Figure 2A). Importantly, the simultaneous knockdown of KDM7A and UTX down-regulated a broad range of NF-κB-dependent genes (Figure S2E-K and Table S2). A principal component analysis (PCA) revealed that the expression patterns of BAY 11-7082- and siKDM7A+siUTX-treated ECs were similar (Figure 2B). Collectively, these transcriptome analyses demonstrated that KDM7A and UTX play critical roles in the TNF-α-induced NF-κB pathways independently, but have a partially synergistic effect in inflammatory ECs.

**Figure 2.**
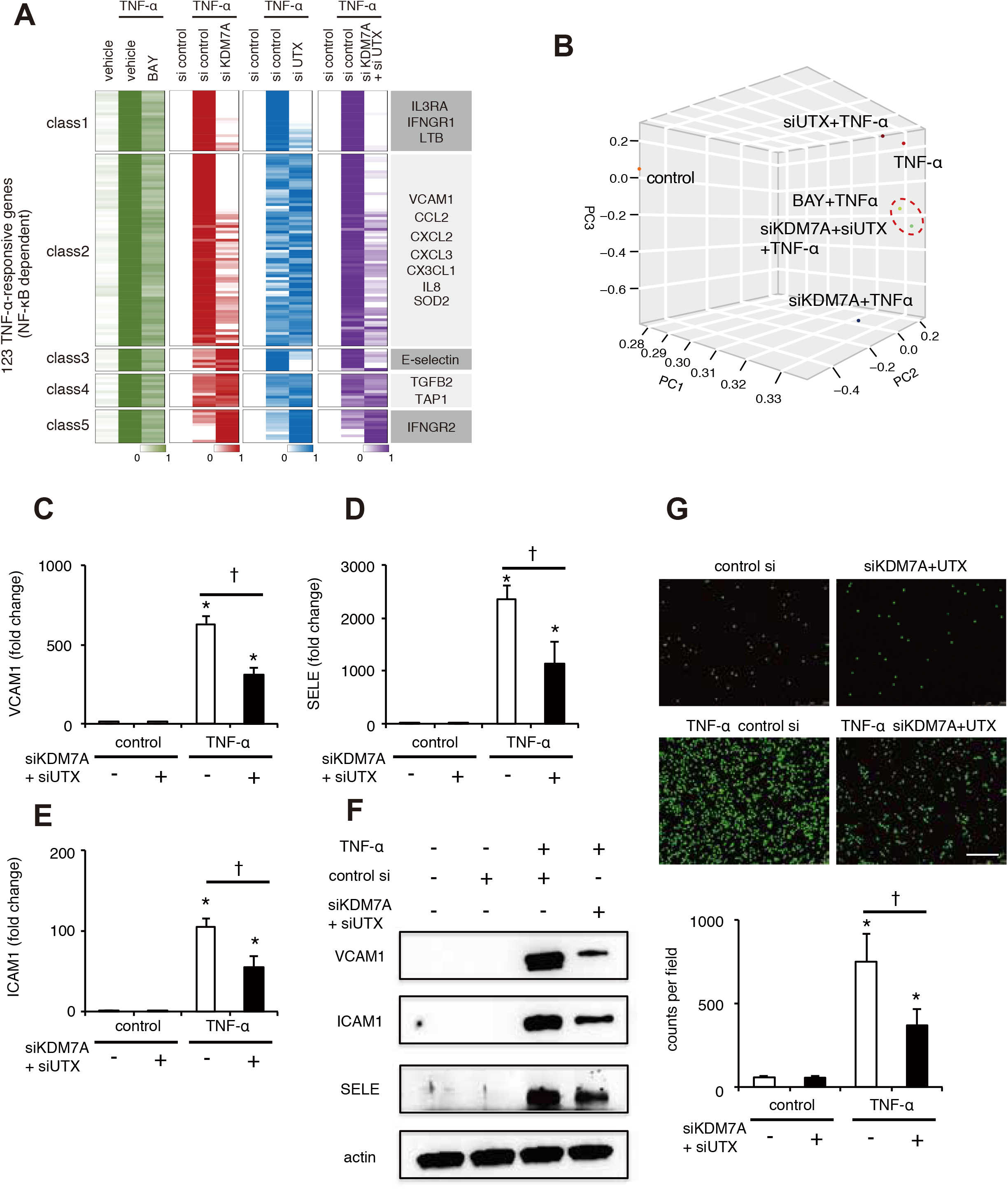
KDM7A and UTX participate in TNF-α-induced NF-κB-p65 signaling pathways leading to the activation of ECs. (A) Heat map representation of RNA-seq results. The p65-dependent genes (total 123; defined as genes up-regulated more than 2-fold via TNF-α treatment and down-regulated more than 0.75-fold via BAY 11-7082 treatment) were classified based on the alteration patterns by the siRNA knockdown of KDM7A, UTX, and both, using the algorithm “HOPACH”. (B) Three-dimensional (3D) multidimensional scaling (MDS) plots. ECs in the presence or absence of TNF-α ± BAY, siKDM7A, siUTX, or siKDM7A+siUTX were mapped to the 3D space, based on the expression profiles of the TNF-α-responsive genes, by using classical MDS. Each treated cell is represented by a colored dot. (C-E) Bar plot showing mean mRNA levels of VCAM1, ICAM1, and SELE measured 4 hrs after stimulation of ECs with or without TNF-α ± (siKDM7A+siUTX). Data are shown as means ± SE. ^*^*P* < 0.05 compared to TNF-α (-) siKDM7A+siUTX (-). † *P* < 0.05 compared to TNF-α (+) siKDM7A+siUTX (-). (F) Western blots for VCAM1, ICAM, SELE, and β-actin in lysates from ECs treated with or without TNF-α (4 hrs) ± (siKDM7A+siUTX). (G) Representative images (top) and bar plot quantification (bottom) showing the adhesion of calcein-labeled U937 monocytes to ECs treated with or without TNF-α ± (siKDM7A+siUTX). Scale bar represents 250 μm. Data are shown as means ± SD. ^*^*P* < 0.05 compared to TNF-α (-) siKDM7A+siUTX (-). † *P* < 0.05 compared to TNF-α (+) siKDM7A+siUTX (-).

We next investigated whether KDM7A and UTX are actually involved in the monocyte adhesion of TNF-α-activated ECs. Consistent with the RNA-seq results (Figure 2A), the siKDM7A treatment significantly decreased the TNF-α-induced expression of VCAM1, but not SELE (Figure S3A-E). In contrast, the knockdown of UTX significantly decreased the TNF-α-induced expression of SELE, but not VCAM1 (Figure S3G-K). Notably, the simultaneous knockdown of KDM7A and UTX resulted in a significant reduction in the TNF-α-induced expression of VCAM1, ICAM1, and SELE, which led to the inhibition of monocyte adhesion to ECs (Figure 2C-G, Figure S3F and L). To ensure that the inhibitory effect of adhesion molecule gene expression by the KDM7A- and UTX-knockdown was not a consequence of the transfection itself or off-target effects, we tested another set of siKDM7A and siUTX oligos and confirmed the reproducible results (data not shown). Together, these data suggested that the miR-3679-5p target genes KDM7A and UTX are involved in monocyte adhesion to TNF-α-activated ECs by controlling the expression of adhesion molecules, VCAM1, ICAM1, and SELE.

### KDM7A- and UTX-recruitment coincide with the NF-κB-p65 related region

Since KDM7A and UTX demethylate H3K9me2 and H3K27me3, respectively, these two factors epigenetically activate their binding loci (Lan et al., 2007; Tsukada et al., 2010). Given that ablated KDM7A- and UTX-expression caused transcriptional repression of inflammation related genes, including adhesion molecules (Figure 2A), we hypothesized that these loci were epigenetically repressed and rapidly activated after TNF-α-stimuli. To test this hypothesis, we performed chromatin immunoprecipitation followed by NGS (ChIP-seq) experiments in replicates, to determine the KDM7A and UTX binding sites before and 1 hr after TNF-α stimulation. Since we could not obtain good antibodies against KDM7A and UTX for ChIP, we overexpressed FLAG-tagged KDM7A and UTX, and immunoprecipitated the proteins with anti-FLAG antibodies (Figure S4A). Examples of biological replicates and correlations are depicted in Figure S4B and C. To correlate KDM7A and UTX binding with epigenetic modifications and transcription factor occupancy, we also measured H3K27ac and NF-κB-p65 at the baseline and after 1 hr of TNF-α stimulation. As a result, the TNF-α-inflammatory signal caused rapid and global redistribution of KDM7A, from 1,714 to 2,483 binding regions. Similarly, we identified 1,303 and 933 UTX associated regions in resting and inflammatory ECs, respectively. In TNF-α-stimulated ECs, the enrichment of KDM7A, UTX, and p65 was evident at promoters (KDM7A: 30.2%, UTX: 20.1%, p65: 20.1%), intragenic regions (KDM7A: 49.6%, UTX: 43.9%, p65: 47.3%), and intergenic regions (KDM7A: 20.2%, UTX: 36.0%, p65: 32.6%) (Figure S4D and H). In response to TNF-α, KDM7A- and UTX-binding was increased in many TNF-α-up-regulated genes, including important adhesion molecules and chemokines, as confirmed by an integrative analysis combined with the RNA-seq and ChIP-seq signal ratios (Figure 3A and B). In accordance with the RNA-seq results (Figure 2A and B), KDM7A bound a larger number of TNF-α-up-regulated genes than UTX (KDM7A: 66, UTX: 33). In contrast, KDM7A- and UTX-binding was decreased in TNF-α-down-regulated genes mainly related to developmental pathways (Figure 3A and B). Consistently, an analysis with the Genomic Regions Enrichment of Annotations Tool (GREAT) demonstrated that the regions with gains of KDM7A or UTX peaks in TNF-α-stimulated ECs showed enrichment for inflammatory pathways including NF-κB and TNF receptor signaling, while the regions with a loss of their peaks showed enrichment for important developmental pathways, including Notch and Wnt signaling (Table S3). For example, the upper right parts of Figure 3A and 3B include VCAM1, SELE, IL8, and CCL2. In contrast, the lower left parts include SOX18 and BMP4. At the coding gene locations, KDM7A and UTX occupancies around the gene body were increased in TNF-α-upregulated genes, and decreased in TNF-α-downregulated genes after TNF-α treatment (Figure 3C and D, and Figure S4E-G). These results demonstrated that the genes with expression that is either up- or down-regulated by TNF-α signaling showed increased and decreased occupancies of the two histone modification enzymes, respectively.

**Figure 3.**
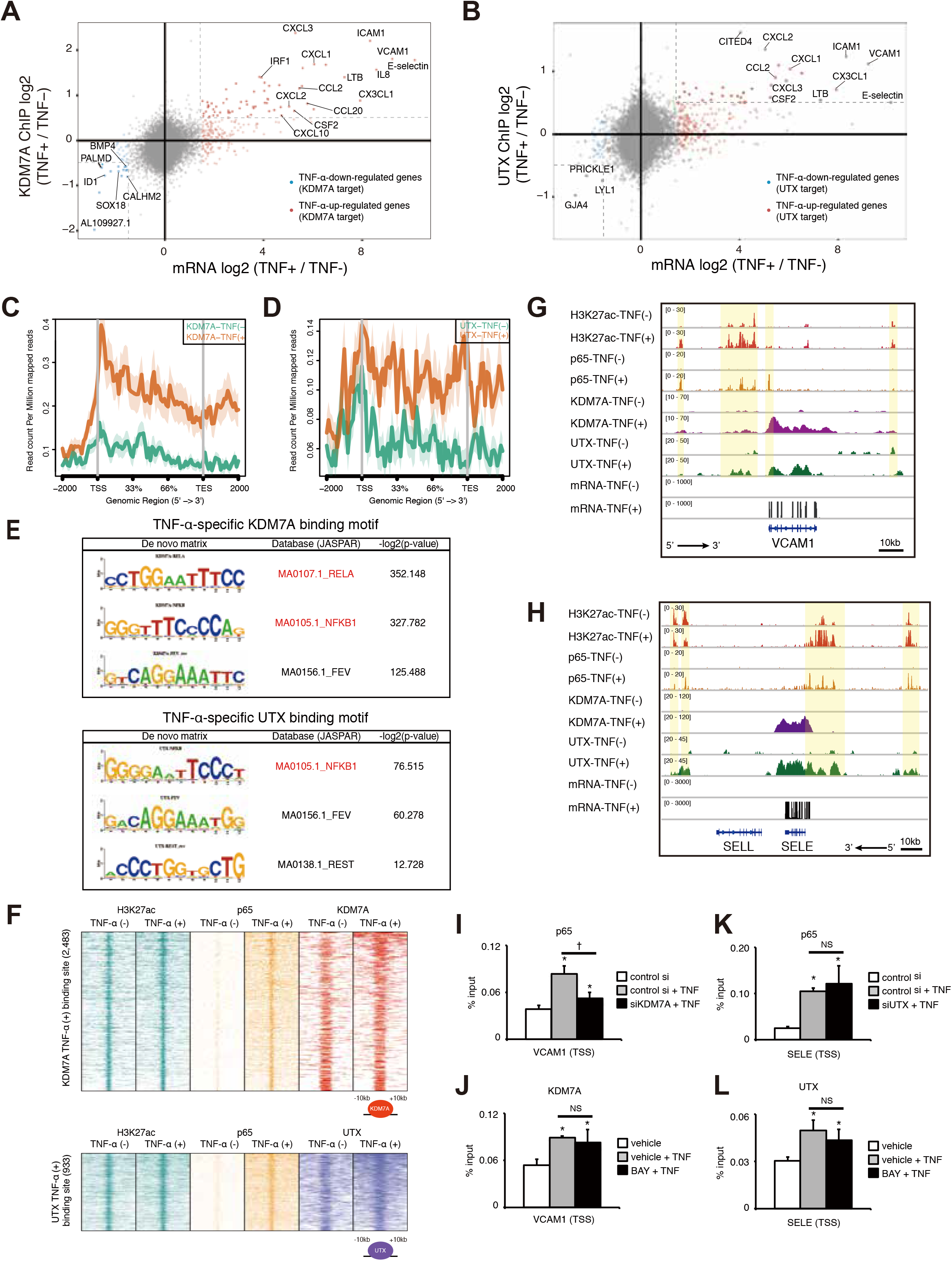
ChIP-seq revealing the recruitment of KDM7A and UTX to NF-κB-p65 related elements. (A and B) Scatter plots showing the connection between KDM7A or UTX binding and RNA expression in TNF-α-stimulated ECs. The X and Y axes indicate the log2 ratio of fragments per kilobase of exon per million mapped sequence reads (FPKM) values and the ChIP-Seq signals (TNF-α 60min/0min) for KDM7A (A) and UTX (B), respectively. Genes with more than a 1.5-fold increase in FPKM values (light red dots) satisfying a 0.5-fold increase in ChIP-seq signals are indicated as dark red dots. Genes with a 1.5-fold decrease in FPKM values (light blue dots) satisfying a 0.5-fold decrease in ChIP-Seq signals are indicated as dark blue dots. These genes are defined as the target genes for KDM7A (A) and UTX (B), respectively. The names of representative genes are shown in the graphs. (C and D) Metagene representations of KDM7A (C) and UTX (D) ChIP-seq signals in units of read count per million mapped reads at a meta composite of the genomic regions around the TNF-α-up-regulated genes. (E) *De novo* motif analyses for KDM7A (top) and UTX (bottom) binding recognition sequences in TNF-α-stimulated ECs. The *P*-value indicates significant enrichment of the transcription factor binding motif in KDM7A/UTX binding sites, in comparison to a size-matched random background. (F) Heat map of H3K27ac (green), p65 (yellow), KDM7A (red), and UTX (purple) levels in resting and TNF-α-stimulated ECs. Each row shows ± 10 kb centered on the KDM7A (top) or UTX (bottom) peak. Rows are ordered by max KDM7A or UTX signal in each region. The ChIP-seq signal is depicted by color scaled intensities. (G and H) Gene tracks of ChIP-seq signals for H3K27Ac, p65, KDM7A, and UTX, and RNA-seq signals around the *VCAM1* (G) and *SELE* (H) loci in TNF-α (-) and TNF-α (+) ECs. ChIP-Seq and RNA-seq signals are visualized by Integrated Genome Viewer (http://software.broadinstitute.org/software/igv/). (I and J) ChIP-qPCR of p65 (I) and KDM7A (J) at the *VCAM1* TSS, normalized to input. Data are shown as means ± SD. ^*^*P* < 0.05 compared to TNF-α (-). † *P* < 0.05 compared to TNF-α (+) siKDM7A (-). (K and L) ChIP-qPCR of p65 (K) and UTX (L) at the *SELE* TSS, normalized to input. Data are shown as means ± SD. ^*^*P* < 0.05 compared to TNF-α (-).

To further examine the characteristics of the KDM7A- and UTX-binding sites, we performed a motif analysis and found that RELA (NF-κB-p65) and NFKB1 (NF-κB-p105) binding sequences were significantly enriched, as compared with random genomic background sequences (Figure 3E). The co-localization of KDM7A/UTX and p65 was also confirmed by a global enrichment analysis (Figure 3F). From the perspective of p65 binding sites, we identified the recruitment of substantial levels of KDM7A and UTX (Figure S4I and J). At the *VCAM1* locus, TNF-α stimulation of ECs for 1 hr increased p65, KDM7A, and UTX occupancy at promoters and surrounding enhancer-like elements marked by H3K27ac (Figure 3G). Coincident with these observations, comparable evidence was obtained at the *SELE* locus, where TNF-α treatment recruited both KDM7A and UTX to the promoter and enhancer-like elements co-localized with p65 (Figure 3H). It should be noted that KDM7A and UTX were bound to the gene bodies of *VCAM1* and *SELE*. Taken together, these results suggested that KDM7A and UTX are recruited to NF-κB-p65 related regions and contribute to transcriptional control through TNF-α signaling in ECs.

Although the interplay between epigenetic mediators and transcription factors is crucial for cytokine signal transduction, it is unclear which factor works upstream from the other. Previous studies hinted that the interplay might depend on the cellular environment or the nuclear architecture (Heinz et al., 2015). To examine the temporal relationships between p65 and KDM7A/UTX recruitment to regulatory elements in this context, we performed siRNA experiments for KDM7A and UTX, followed by ChIP-qPCR for p65 enrichment. As shown in Figure 3I, the knockdown of KDM7A reduced the TNF-α-induced enrichment of p65 at the *VCAM1* transcription start site (TSS). However, the BAY 11-7082 treatment did not affect the recruitment of KDM7A at the *VCAM1* TSS (Figure 3J). At the *SELE* TSS, the knockdown of UTX and the BAY 11-7082 treatment had no effect on the recruitment of p65 and UTX, respectively (Figure 3K and L). Thus, KDM7A, but not UTX, might be required for the TNF-α-induced recruitment of p65 to regulatory elements in ECs.

### TNF-α immediately removes H3K9me2 and H3K27me3 marks around KDM7A- and UTX-target genes

The KDM7A- and UTX-mediated demethylation of repressive histone marks (H3K9me2 and H3K27me3, respectively) correlates with active gene expression (Agger et al., 2007; Lee et al., 2007; Tsukada et al., 2010). Therefore, we hypothesized that the transcriptional activation of genes in TNF-α-stimulated ECs would involve KDM7A- and UTX-dependent histone modifications. To address this hypothesis, we examined the epigenetic landscape for H3K9me2 and H3K27me3 in ECs after TNF-α signaling by ChIP-seq. As shown in Figure 4A and B, the ChIP-seq signals for H3K9me2 and H3K27me3 around important adhesion molecules and chemokines were immediately decreased in response to TNF-α treatment, although the global levels of H3K9me2 and H3K27me3 around KDM7A- and UTX-target genes remain unchanged. At the *VCAM1* locus, the ChIP-seq signals for H3K9me2 were broadly decreased 1 hr after TNF-α treatment, which coincides with KDM7A recruitment (Figure 4C). ChIP followed by qPCR also revealed a significant reduction in H3K9me2 levels around the TSS and enhancer regions of *VCAM1* (Figure 4D). There was no obvious reduction in ChIP-seq signals for H3K27me3 around the *VCAM1* locus, while ChIP followed by qPCR detected a significant reduction especially within the enhancer regions, but not the TSS (Figure 4C and E). Similarly, at the *SELE* locus, the H3K9me2 and H3K27me3 levels detected by ChIP-seq and ChIP-qPCR were broadly decreased in association with KDM7A- and UTX-recruitment in both the TSS and enhancer regions (Figure 4F-H). In contrast, TNF-α treatment did not affect the H3K9me2 and H3K27me3 levels at the genomic regions of *HOXA13* and *GATA4*, which are not or only weakly expressed in human ECs with or without an exogenous stimulus (Figure 4I and J) (Kanki et al., 2011). Next, to test whether the repressive histone mark reduction rapidly induced by TNF-α stimulation was directly due to the two histone modifiers, we further assessed the influence of the knockdowns of KDM7A and UTX on the H3K9me2 and H3K27me3 levels in the genomic regions around *VCAM1* and *SELE*, respectively. The siRNA knockdowns of KDM7A and UTX restored H3K9me2 and H3K27me3 at representative regions of *VCAM1* and *SELE*, respectively, in TNF-α-treated human ECs (Figure 4K and L). Taken together, these results suggested that the KDM7A- and UTX-mediated regulation of repressive histone marks might be a key mechanism for the transcriptional activation of inflammatory genes in human ECs.

**Figure 4.**
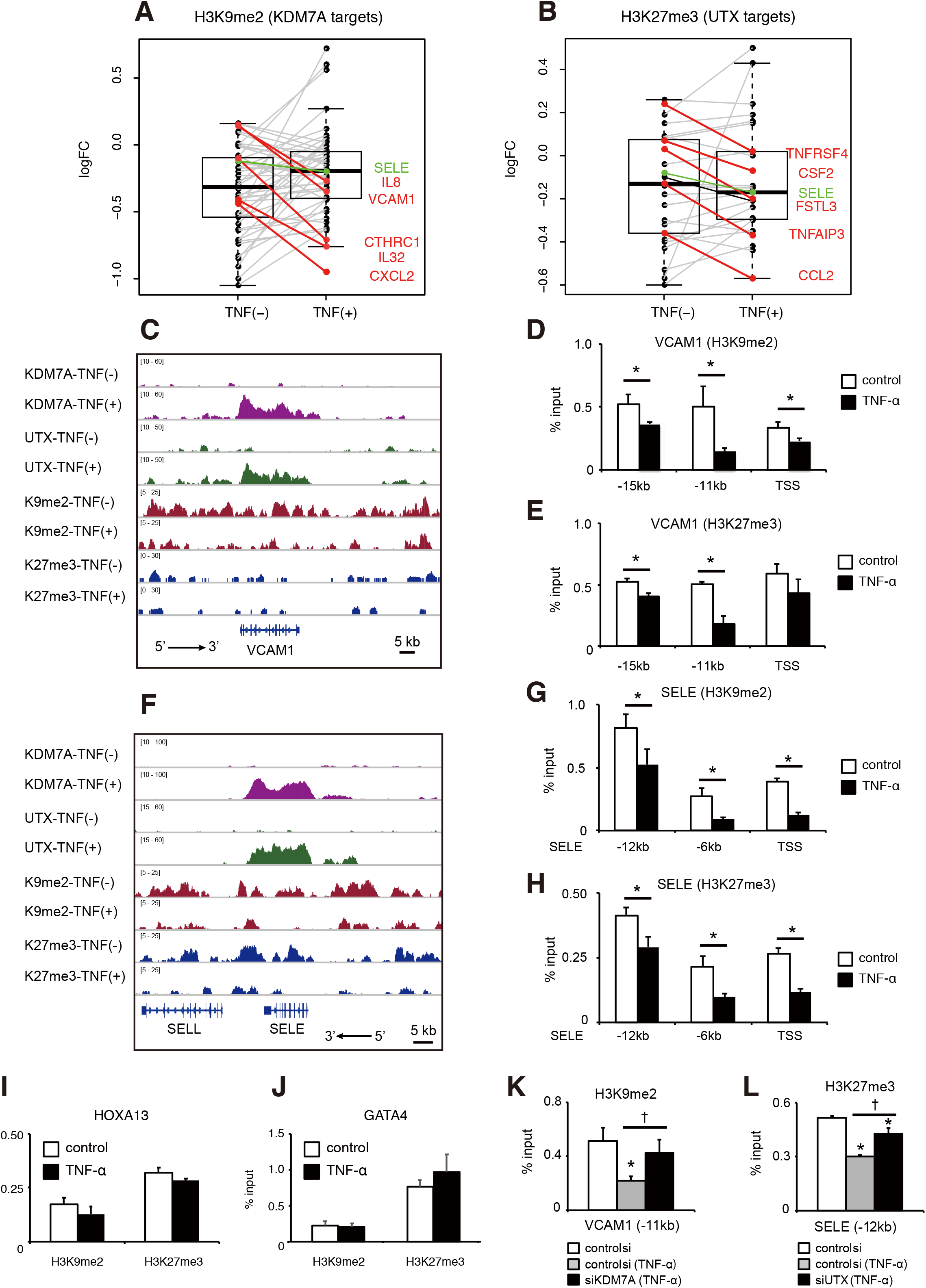
H3K9me2 and H3K27me3 marks around KDM7A- and UTX-target genes change after TNF-α-treatment. (A) Box plot of the log2 fold change of ChIP-seq enrichment of H3K9me2, compared to input DNA, in KDM7A target genes with TNF-α treatment (TNF+) and control (TNF-). Each dot in the box plot shows data for one single target gene. Genes colored red represent the top five downregulated target genes with TNF-α-treatment, as compared to control. (B) Box plot of the log2 fold change of ChIP-seq enrichment of H3K27me3, as compared to input DNA, in UTX target genes with TNF-α treatment (TNF+) and control (TNF-). Each dot in the box plot shows data for one single target gene. Genes colored red show the top five downregulated target genes with TNF-α treatment, as compared to control. (C and F) Gene tracks of ChIP-seq signals for KDM7A, UTX, H3K9me2, and H3K27me3 around the *VCAM1* (C) and *SELE* (F) loci in TNF-α (-) and TNF-α (+) ECs. (D and E) ChIP-qPCR showing enrichment (percent input) of H3K9me2 (D) and H3K27me3 (E) around the *VCAM1* locus. Primer pairs targeting distinct regions are listed on the X axis. (G and H) ChIP-qPCR showing enrichment (percent input) of H3K9me2 (G) and H3K27me3 (H) around the *SELE* locus. Primer pairs targeting distinct regions are listed on the X axis. (I and J) ChIP-qPCR showing enrichment (percent input) of H3K9me2 and H3K27me3 around the *HOXA13* (I) and *GATA4* (J) loci. Data are shown as means ± SD. ^*^*P* < 0.05 compared to TNF-α (-). (K and L) ChIP-qPCR showing enrichment (percent input) of H3K9me2 around *VCAM1*-11kb (K) and H3K27me3 around *SELE*-12kb (L). Data are shown as means ± SD. ^*^*P* < 0.05 compared to TNF-α (-). † *P* < 0.05 compared to TNF-α (+) siKDM7A (-) or TNF-α (+) siUTX (-).

### TNF-α-induced SEs correlate with KDM7A and UTX recruitment and chromatin conformation change

SEs are genomic regions consisting of clusters of regulatory elements bound with extremely high amounts of transcription factors, and are associated with characteristic histone modifications and genomic architecture. Recent studies have shown that the interaction between NF-κB-p65 and BRD4 on SEs rewires the gene expression program toward proinflammatory responses in human ECs. We found that the ChIP-seq signals for BRD4 (Brown et al., 2014) and H3K27ac increased at the KDM7A- and UTX-binding sites in TNF-α-treated human ECs (Figure S5A and B). Thus, we hypothesized that the recruitment of KDM7A and UTX was coincident with and functionally associated with TNF-α-induced SEs. To test this hypothesis, we used ROSE (Whyte et al., 2013) and defined the SEs in human ECs based on the ChIP-seq signals for BRD4 (Brown et al., 2014). When ranked by increasing BRD4 enrichment, 311 and 356 SEs were identified in resting (TNF-α-untreated) and TNF-α-treated ECs, respectively. Many of these SEs are identical to the KDM7A- and UTX-bound regions (Figure 5A and Figure S5C). These identical SEs in TNF-α-treated ECs were located adjacent to inflammatory gene loci, including *VCAM1*, *IL8*, *SELE*, and *CCL2*, while those in resting ECs were located adjacent to gene loci critical to non-inflammatory EC functions, such as *SOX18*, *NR2F2*, *ESAM*, and *PECAM1*. Furthermore, as compared to typical enhancers (TEs: obtained from the pre-compiled HUVEC ngsplotdb), KDM7A and UTX binding was dramatically increased at SEs (Figure 5B and C). These results indicated the possible roles of KDM7A and UTX in the function or formation of SEs in human ECs.

**Figure 5.**
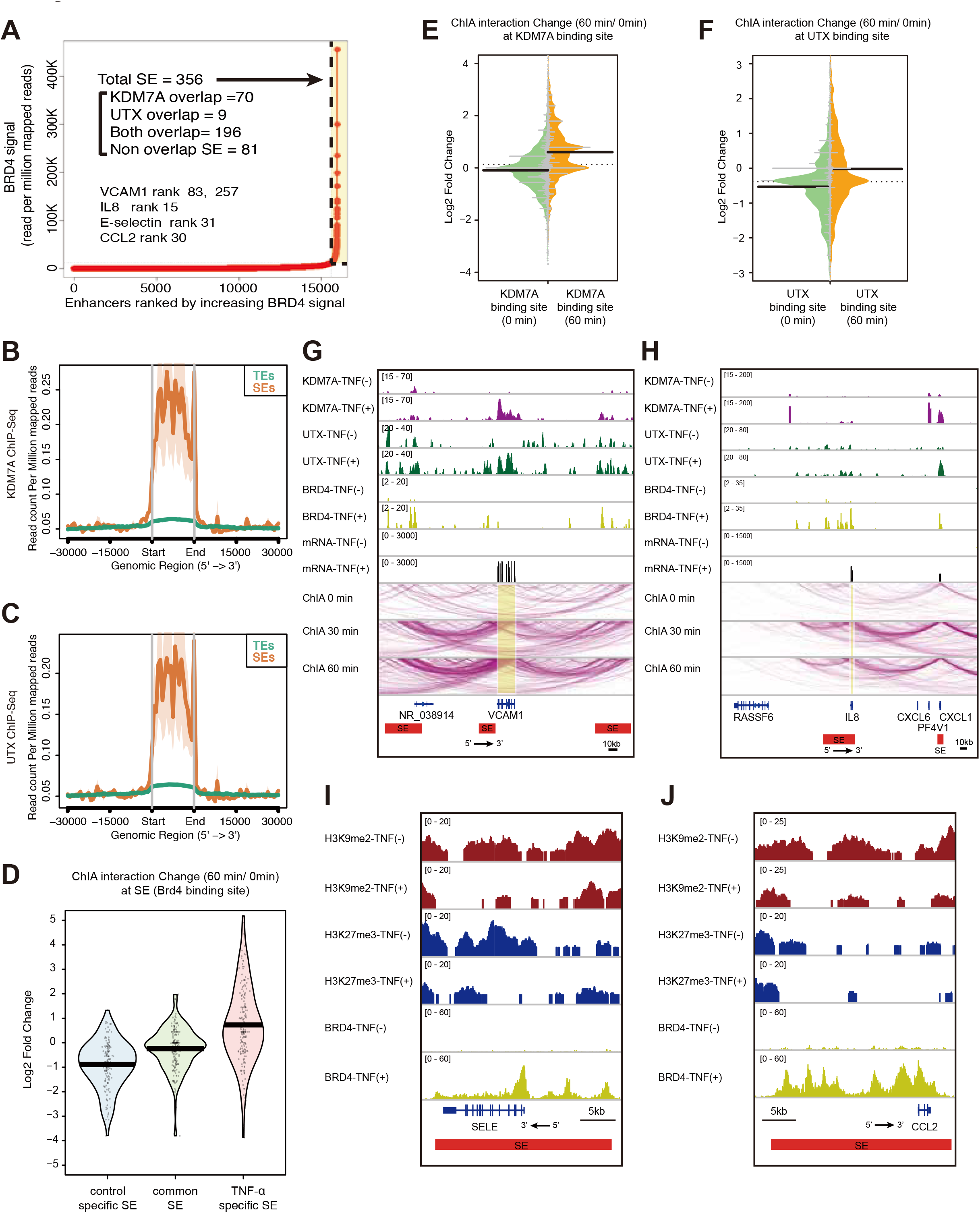
The recruitment of KDM7A and UTX is associated with the formation of TNF-α-induced SEs. (A) Plot of enhancers defined in TNF-α-treated ECs, ranked by increasing bromodomain-containing protein 4 (BRD4) signal. Super enhancers (SEs) were defined by ROSE (Loven et al., 2013; Whyte et al., 2013), based on the ChIP-Seq signals for BRD4 (Brown et al., 2014). SEs are indicated by dashed lines and are colored yellow. The numbers show the total SEs and their classification based on KDM7A and UTX binding. Representative SE-related genes are indicated in the graph. (B and C) Metagene representations of KDM7A (B) and UTX (C) ChIP-seq signals in units of read count per million mapped reads, at a meta composite of SEs and typical enhancers (TEs). TEs were obtained from the pre-compiled HUVEC ngsplotdb (https://github.com/shenlab-sinai/ngsplot). (D) Pirate plots showing ChIA-PET interaction changes at control-specific, common, and TNF-α-specific SEs. The Y axis indicates the log2 ratio of ChIA-PET signals (TNF-α 60 min/0 min). The central bar indicates the average. (E and F) Bean plots showing ChIA-PET interaction changes at KDM7A (E) and UTX (F) binding sites. The Y axis indicates the log2 ratio of ChIA-PET signals (TNF-α 60 min/0 min). The central bar indicates the median. (G and H) ChIA-PET interactions of active RNA pol II around the *VCAM1* (G) and *IL8* (H) loci integrated with the ChIP-seq profiles of KDM7A, UTX, and BRD4, and RNA-seq profiles in TNF-α (-) and TNF-α (+) ECs. ChIA-PET interactions were visualized by the WashU Epigenome Browser (http://epigenomegateway.wustl.edu/browser/). Interactions detected by ChIA-PET are depicted with purple lines. Red bars show TNF-α-specific SEs. (I and J) Gene tracks of ChIP-seq signals for H3K9me2, H3K27me3, and BRD4 around *SELE* (I) and *CCL2* (J) in TNF-α (-) and TNF-α (+) ECs. Red bars show TNF-α-specific SEs.

To investigate whether EC SEs can be used to prioritize non-coding functional variants for cardiovascular diseases, we overlapped the physical coordinates of the 356 SEs from Figure 5A with the genome-wide association study (GWAS) associated variants. Single nucleotide polymorphisms (SNPs) satisfying the genome-wide significance for coronary artery disease (CAD), atherosclerosis, hypertension, or stroke were downloaded from the NHGRI-EBI GWAS Catalog (Welter et al., 2014). To account for linkage disequilibrium (LD) between closely spaced SNPs on the same chromosome, we used data from the 1000 Genomes Project (Genomes Project et al., 2015) to retrieve SNPs in LD with the reported GWAS SNPs, when r2 was greater than 0.8 based on European haplotype structure. We identified 29 SNPs that were within EC SEs, in which 12 SNPs overlapped with both the KDM7A- and UTX-binding regions and 9 SNPs overlapped with the KDM7A-binding regions (Table S4). Importantly, other non-overlapping SNPs were also located adjacent to the KDM7A- and UTX-binding regions. These data provided a focused list of potential functional non-coding variants that can affect the susceptibility to cardiovascular diseases through EC gene regulation, and suggested that the predisposition mechanisms of these variants might be associated with KDM7A and UTX functions at SEs.

The relationship between SEs and genomic architecture has been implicated, especially in stem cells and cancer cells, by mapping the local chromosomal structure using ChIA-PET (Dowen et al., 2014; Handoko et al., 2011; Hnisz et al., 2016). To capture the active formation of chromatin loops through TNF-α signaling in human ECs, we performed ChIA-PET using an antibody against the phospho-Ser2/Ser5 heptad repeats in the C-terminus of the largest catalytic subunit of RNA polymerase II (active RNAP). The active RNAP ChIA-PET data identified 5,948,648, 4,849,886 and 3,893,377 interactions at 0, 30, and 60 min after the TNF-α treatment, respectively, suggesting that the active RNAP interactions slightly decreased overall but were concentrated on specific inflammatory response loci. These ChIA-PET data were further processed by the ChIA-PET tools (Li et al., 2010), which identified an average of 4,280 high-confidence interactions (false discovery rate of 1% and *P* < 0.0001), including 159 SE-SE interactions. The SE-SE loops had a median length of 7,023. We next investigated whether the SE-SE interactions could be changed upon TNF-α stimulation in human ECs. We compared the ChIA-PET interaction changes between control-specific, common, and TNF-α-specific SEs. Interestingly, the ChIA-PET interactions increased at TNF-α-specific SEs in response to TNF-α, while they decreased at control-specific SEs (Figure 5D). Importantly, the ChIA-PET interactions increased at both the KDM7A- and UTX-binding sites in TNF-α-treated ECs (Figure 5E and F). As shown in Figure 5G and H, the SE-SE interactions were newly formed within 1 hr after TNF-α treatment at the *VCAM1* and *IL8* loci, and were associated with the recruitment of KDM7A and UTX. In contrast, in response to TNF-α, the ChIA-PET interactions decreased at the loci of genes critical for EC function, such as *SOX18* and *NR2F2*, in conjunction with decreased transcription and KDM7A- and UTX-recruitment (Figure S5D and E). As exemplified by the *SELE* and *CCL2* loci, the H3K9me2 and H3K27me3 levels decreased at SEs in TNF-α-treated human ECs (Figure 5I and J). Collectively, these data suggested that KDM7A and UTX might be functionally involved in the formation of SEs and the chromosomal conformation changes activating their associated genes during early inflammatory responses in human ECs.

### TNF-α immediately induces chromosomal conformation changes within sub-TADs in human ECs

Although the active RNAP ChIA-PET data indicated that chromosomal conformation changes immediately occurred after TNF-α treatment in human ECs, the ChIA-PET analysis was methodologically biased by the factor-dependent immunoprecipitation step and therefore could not directly reflect the chromosomal conformation changes. To address this point, we performed *in situ* Hi-C, which allows the unbiased identification of chromatin interactions across an entire genome. We obtained a total of approximately 200 million reads from two biological replicates under each condition (*i.e*., TNF-α 0 min: replicate 1, replicate 2, and TNF-α 60 min: replicate 1, replicate 2). Since the two biological replicates showed a high degree of correlation (Figure S6A and B), the Hi-C data from the two biological replicates were combined for further analysis. We detected 10,577 and 10,851 interactions in the TNF-α-untreated and -treated human ECs, respectively, among which 7,196 interactions were shared. The Hi-C matrix identified megabase size TADs with boundaries that were highly similar between the TNF-α-untreated and -treated human ECs (Figure 6A and B). Indeed, the total number of TADs, which was calculated by TADtool (Kruse et al., 2016), remained relatively unchanged before (6,349) and after (7,103) TNF-α treatment. Importantly, this is consistent with previous reports showing that megabase size TADs are largely conserved among different tissues, and are already formed in embryonic stem cells and remain relatively constant during development (Dixon et al., 2012).

**Figure 6.**
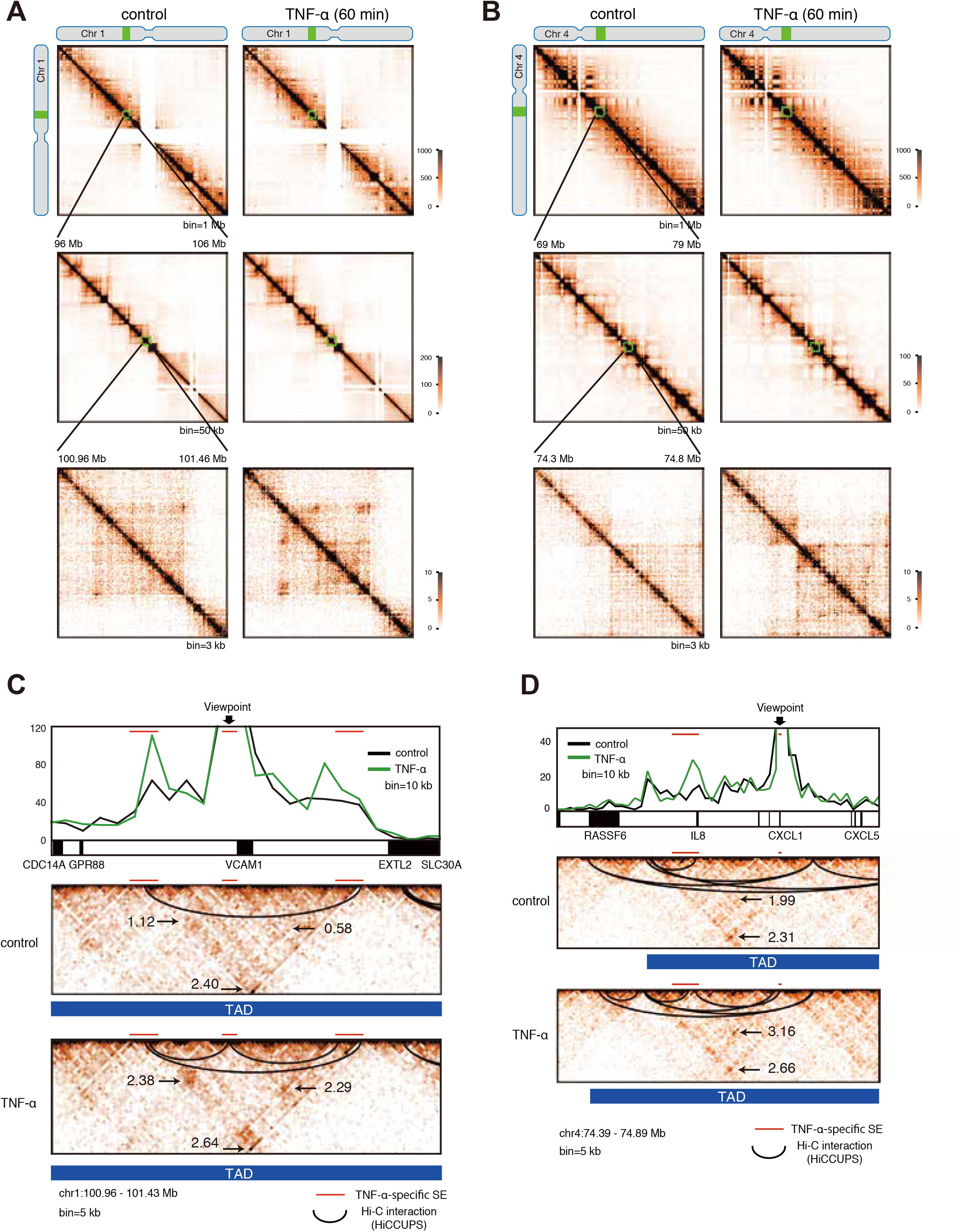
TNF-α treatment rapidly induced chromosomal conformation changes in human ECs. (A and B) Hi-C contact matrices from chromosome 1 (A) and chromosome 4 (B): the whole chromosome, at 1 Mb resolution (top); 50 kb resolution (middle); 3 kb resolution (bottom). Left: TNF-α 0 min; Right TNF-α 60 min. Green boxes indicate regions used for zoom in. The highest resolution maps show 500 kb regions including the *VCAM1* (A) and *IL8* (B) loci, respectively. The color intensity of each pixel represents the normalized number of contacts between a pair of loci. (C and D) Virtual 4C views and heat map of Hi-C data around the *VCAM1* (C) and *IL8* (D) loci. Virtual 4C is calculated from Hi-C datasets, using bins covering the indicated viewpoints. The Y axis in the virtual 4C view indicates the number of reads that interact to the viewpoint bin. Control reads are raw counts, and TNF-α-reads were normalized to control reads. Valid chromatin loops were calculated by HiCCUPS (https://github.com/theaidenlab/juicer/wiki/Download) and are shown with black lines. Topologically associating domains (TADs) were calculated by TADtool (https://github.com/vaquerizaslab/tadtool) and are shown as blue bars. The number denotes the Aggregate Peak Analysis (APA) score at the indicated areas (Rao et al., 2014). Red lines indicate TNF-α-specific SEs.

Since the SEs and their associated genes are located within sub-TAD boundaries (~1 Mb) (Dowen et al., 2014; Tang et al., 2015), we further analyzed the “intra-TAD” interactions occurring within sub-TADs. Virtual 4C, derived from Hi-C data at the representative *VCAM1* and *IL8* loci, showed an increase of several SE-SE interactions within 1 hr after TNF-α treatment in human ECs (Figure 6C and D). Accordingly, HiCCUPS, an algorithm for finding chromatin loops (Rao et al., 2014), identified newly formed SE-SE loops in TNF-α-treated human ECs around the genomic regions of *VCAM1* (2 loops) and *IL8* (1 loop) (Figure 6C and 6D). Importantly, these newly formed SE-SE loops were located within sub-TADs, which were similar before and after the TNF-α treatment in human ECs. In addition, the Aggregate Peak Analysis (APA) (Rao et al., 2014) revealed that the loops only found in TNF-α-treated ECs were strengthened by TNF-α signaling (1.12 to 2.38 or 0.58 to 2.29 at *VCAM1* SEs, 1.99 to 3.16 at *IL8* SEs) (Figure 6C and D). In contrast, the Hi-C interactions were unchanged at the genomic regions of the *PALMD* and *HOXA* loci, which exist beside the control-specific and common SEs, respectively (Figure S6C and S6D). Taken together, these data indicated that the chromosomal conformation changes within sub-TADs could occur at the active transcriptional regions during the early inflammatory response in human ECs.

### KDM7A and UTX are important for leukocyte adhesion in mice

Recent studies demonstrated the utilities of the KDM2/7 subfamily inhibitor Daminozide and the KDM6 subfamily inhibitor GSK-J4 for treating breast cancer and brainstem glioma, respectively (Chen et al., 2016; Hashizume et al., 2014). To explore the phenotypic effects of the pharmacological inhibition of KDM7A and UTX, we tested the functional effects of Daminozide and GSK-J4 on leukocyte rolling across TNF-α-activated endothelium *in vivo*. C57BL/6 mice were pretreated with Daminozide alone (50 mg/kg), GSK-J4 alone (50 mg/kg), or Daminozide in combination with GSK-J4 (50 mg/kg each) at 16 hrs and 1 hr prior to TNF-α-injection (Figure 7A). The leukocyte adhesive interaction in the left femoral artery was observed by intravital microscopy, which showed that Daminozide in combination with GSK-J4 significantly decreased the number of interacting leukocytes induced by TNF-α-injection (36.7 versus 8.4, p < 0.05; Figure 7B and C). Importantly, Daminozide alone and GSK-J4 alone did not improve the leukocyte adhesive interactions (Figure S7A and B), suggesting that KDM7A and UTX are cooperatively and synergistically involved in endothelium activation in mice.

**Figure 7.**
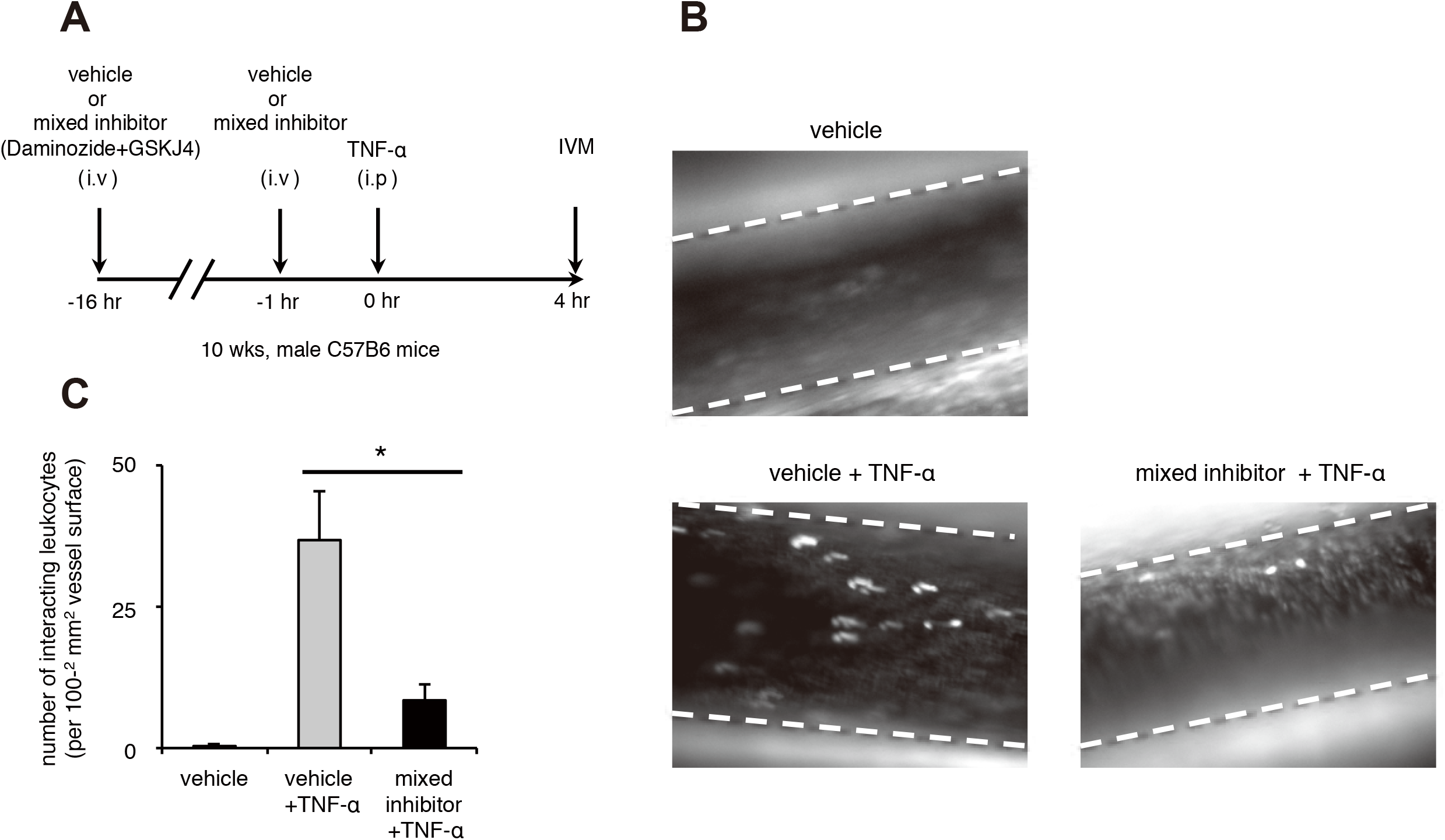
KDM7A and UTX are important for leukocyte adhesion in mice. (A) Drug administration protocol for intravital microscopy (IVM) experiments. Male, 10 week old C57BL/6 mice (n=3/group) were pretreated with vehicle or mixed inhibitor (Daminiozed+GSKJ4, 50 mg/kg each) at 16 hrs and 1 hr prior to TNF-α treatment. Inflammation was induced by an intraperitoneal injection of TNF-α (5 μg/mouse) and IVM was performed 4 hrs later. (B) Representative snapshot from the IVM analysis of leukocyte adhesive interactions in the femoral arteries. Margins of vessels are indicated with dashed lines. White spots represent fluorescently labeled leukocytes visualized by the intravenous injection of rhodamine 6G. (C) Quantitative analyses of leukocyte adhesive interactions in the femoral arteries. Data are shown as means ± SE. ^*^*P* < 0.05 compared to the vehicle group. † *P* < 0.05 compared to the vehicle + TNF-α group. See also Movies S1 and 2.

### Discussion

The inflammatory response plays an important role in tissue homeostasis and its dysregulation underlies numerous diseases (Gistera and Hansson, 2017; Libby, 2002; Libby et al., 2011), and thus it is crucial to understand how these processes are regulated. The master transcription factor, NF-κB, directly controls the induction of a very large fraction of inflammatory genes (Zhang et al., 2017). Recent genome-wide studies in differentiated cells have revealed the essential role of epigenetic regulation in orchestrating the information from an inflammatory cascade to regulate NF-κB recruitment to specific genomic sites and activate NF-κB-dependent gene expression programs. In mouse macrophages, most enhancers that are bound by NF-κB following an inflammatory stimulus are already pre-bound by the hematopoietic lineage factor PU.1 (Ghisletti et al., 2010; Link et al., 2018), while the *de novo* formation of latent NF-κB enhancers depends on SWI/SNF chromatin-remodeling complexes being occupied by PU.1 and associated transcription factors (Ostuni et al., 2013). Consistently, about half of the NF-κB-bound enhancers are pre-bound by both the ETS and AP1 transcription factors in human ECs (Hogan et al., 2017). On the other hand, the activation of human ECs with TNF-α results in the formation of large NF-κB-bound enhancer clusters (*i.e*., SEs), which are highly occupied by BRD4 and a driving force for activating proinflammatory gene expression (Brown et al., 2014). However, the current comprehension of the epigenetic mechanisms that govern transcriptional activation during early inflammatory responses is far from complete. Here we demonstrated the intrinsic and synergistic roles of two histone demethylases, KDM7A and UTX, in regulating the epigenetic landscape and the inflammatory gene expression program through the TNF-α/NF-κB signaling pathway in human ECs.

Repressive histone marks, H3K9me2 and H3K27me3, have been implicated in transcriptional repression in immune cells. An early study showed the rapid demethylation of H3K9 (the antibody used in this study did not seem to discern the mono-, di-, or tri-methyl state of H3K9) at promoters of activated inflammatory genes in macrophages treated with lipopolysaccharide (LPS) (Saccani and Natoli, 2002). Other work demonstrated the important role for H3K9me2 in the regulation of cytokine expression, which was confirmed by findings of lower levels of H3K9me2 at inflammatory gene promoters in innate immune cells as compared with nonimmune cells, such as mouse embryonic fibroblasts and cardiomyocytes. Genetic deletion and pharmacological inhibition of the H3K9 methyltransferase G9a resulted in the phenotypic conversion of fibroblasts into highly potent cytokine producing cells (Fang et al., 2012). It was also reported that the expression levels of the H3K27 demethylase JMJD3 were increased in mouse macrophages dependent on NF-κB, in response to LPS (De Santa et al., 2007). GSK-J4, a potent enzyme inhibitor of JMJD3 and UTX, reduced LPS-induced proinflammatory cytokine production in human primary macrophages, and this was associated with the retention of H3K27me3 at inflammatory gene loci (Kruidenier et al., 2012). In this study, we found that KDM7A and UTX regulate the expression of many NF-κB-dependent genes in TNF-α-treated human ECs (Figure 2). KDM7A and UTX were rapidly recruited to the loci of inflammatory genes in human ECs in response to TNF-α, and this was highly correlated with NF-κB binding (Figure 3). We also observed that TNF-α stimulation decreased the H3K9me2 and H3K27me3 levels at the target gene loci of KDM7A and UTX, respectively. As typical examples, the H3K9me2 levels at the *VCAM1* locus and the H3K27me3 levels at the *SELE* locus were retained by the knockdowns of KDM7A and UTX, respectively, in TNF-α-treated human ECs (Figure 4J and K). Taken together, our data have identified KDM7A and UTX as previously unknown histone lysine demethylases that influence the transcription of inflammatory genes through TNF-α signaling in human ECs.

Our results have demonstrated that KDM7A and UTX control large and small numbers of NF-κB dependent genes, respectively, in TNF-α-treated human ECs (Figure 2A). This is consistent with the results that KDM7A is recruited to many more TNF-α-induced gene loci than UTX (Figure 3A and B). Thus, in comparison with UTX, KDM7A seems to play a dominant role for regulating inflammatory responses in human ECs. Importantly, some inflammatory genes, including ICAM1, might require both KDM7A and UTX *in vitro*. Indeed, *in vivo*, the inhibition of leukocyte adhesion to the femoral artery in mice was only observed after a treatment with both Daminozide and GSK-J4 (Figure 7A-C and Figure S7A-B). These results suggested that a synergism exists between the two histone demethylases, KDM7A and UTX, in inflammatory activated ECs. Although direct evidence of a functional interaction between H3K9me2 and H3K27me3 during the inflammatory response is still lacking, a subset of developmental genes may be cooperatively repressed by both H3K9me2 and H3K27me3 during ESC differentiation (Kurimoto et al., 2015; Mozzetta et al., 2014). An immunohistochemical analysis of carotid arteries from hypercholesterolemic mice showed global reductions of the H3K9me3 and H3K27me3 levels in vascular smooth muscle cells, but no changes in their levels in ECs (Alkemade et al., 2010). Decreased levels of H3K27me3 were also observed in vascular smooth muscle cells from advanced human atherosclerotic plaques (Wierda et al., 2015). In addition, a ChIP analysis revealed that the H3K9me3 levels at the promoters of important inflammatory genes were decreased in cultured vascular smooth muscle cells derived from diabetic mice, in association with the reduction of the H3K9me3 methyltransferase Suv39h1 (Villeneuve et al., 2008). Thus, the reduction of repressive histone marks may play an important role in inflammatory diseases, including atherosclerosis, but further studies are needed to understand these aspects.

SEs are large genomic regions, with a median size generally an order of magnitude larger than that of normal enhancers, and are occupied by an extremely high density of enhancer-associated factors, including transcription factors, Mediators, histone acethyltransferases (*e.g*., p300/CBP), histone modifications (*e.g*., H3K27ac), RNAP, and non-coding RNAs. SEs normally regulate the expression of genes that have especially prominent roles in cell type-specific processes (Hnisz et al., 2017; Pott and Lieb, 2015). Although numerous studies have reported the importance of SEs in the maintenance of cell identity and the determination of cell lineage, the significance of *de novo* SE formation during inflammatory responses has also been reported in LPS-stimulated mouse macrophages and TNF-α-stimulated human adipocytes (Hah et al., 2015; Schmidt et al., 2015). In human ECs, *de novo* SEs are rapidly formed as a mechanism by which a master transcription factor could connect a rapid transcriptional response that drives dynamic change following proinflammatory activation. NF-κB engaged most endothelial enhancers in response to TNF-α-stimuli, and directed BRD4 recruitment to form *de novo* SEs and activated the transcription of many canonical proinflammatory genes (Brown et al., 2014). Our present findings show that the recruitment of KDM7A and UTX was coincident with NF-κB binding and H3K27ac marks in TNF-α-stimulated human ECs. We defined the SEs by using the ChIP-Seq data for BRD4, and demonstrated that the binding of KDM7A and UTX was dramatically increased at BRD4-SEs, as compared to typical enhancers. In addition, we observed that the KDM7A recruitment preceded the NF-κB binding to the genome (Figure 3I and J). Thus, KDM7A and UTX might play important roles for the formation of *de novo* SEs in human ECs, although loss of function experiments will be required to address this point.

A recent study described the potential mechanisms by which the repressive histone mark H3K27me3 controls the formation of SEs. SEs identified in hair follicle stem cells and their differentiated progeny were either repressed or activated in a lineage-dependent manner. SEs and their surrounding genomic regions associated with genes critical for determining stem cell fate were actively repressed by gaining H3K27me3 marks, in a process termed “super-silencing” during hair follicle stem cell differentiation. Conversely, differentiated progeny fate determinants became de-repressed (activated) by losing H3K27me3 marks to expose new SEs (Adam et al., 2015). Here, in human ECs, we have shown that KDM7A and UTX rapidly mobilized to newly formed SEs where the repressive histone marks H3K9me2 and H3K27me3 were decreased 1 hr following the TNF-α stimulus, and were associated with the active transcription of many proinflammatory genes (Figure 5I and J). In contrast, KDM7A and UTX were de-localized from the control-specific SEs (TNF-α-lost SEs) in response to TNF-α, despite the persistent hypomethylation of H3K9me2 and H3K27me3 (data not shown). Thus, these findings provide evidence that repressive histone marks might play important roles for both SE-formation and SE-loss during the early inflammatory responses in completely differentiated cells. It was recently demonstrated that heterochromatic domain formation (HP-1-H3K9me2/3 binding) could be rapidly driven by phase separation, a phenomenon that gives rise to diverse, non-membrane-bound compartments (Larson et al., 2017; Strom et al., 2017). The phase separation model could explain the formation of SEs (Hnisz et al., 2017; Sabari et al., 2018). Thus, the heterochromatic states formed by phase separation might involve the formation or loss of SEs, although further studies are needed to investigate whether these models can actually be applied to SE formation.

3C-based technologies have revealed that megabase size TADs are already formed in ES cells and remain relatively constant throughout development (Dixon et al., 2015; Dixon et al., 2012). However, chromosome reorganization is observed during cellular differentiation at sub-TAD levels (*i.e*., within megabase size TADs), and these lineage-specific sub-TADs are associated with cell type-specific gene expression programs (Phillips-Cremins et al., 2013). However, only few studies have investigated the plasticity of chromosome organization or chromosomal loop formation in response to external stimuli. For example, using Hi-C and 3C methods, a report has confirmed that promoter-enhancer loops pre-exist and are relatively stable during TNF-α signaling in human ECs (Jin et al., 2013). In this study, we performed *in situ* Hi-C in combination with active RNA pol II ChIA-PET, and found that the megabase size TADs are highly similar between untreated and TNF-α-treated human ECs. However, to our surprise, SE-SE interactions are newly formed at sub-TAD levels at typical TNF-α-responsive inflammatory gene loci, 1 hr after TNF-α stimulation. There are two possible explanations for this phenomenon. First, the differences in the restriction enzymes used in each study might be the reason for this observation. We used the 4-cutter restriction enzyme *Mbo*l, whereas the other study used the 6-cutter restriction enzyme *Hin*dIII. *Mbo*I targets 7,227,513 sites in the human genome, which is about ten-fold more than *Hin*dIII (846,132), and thus could theoretically capture more chromosome contact information (Lin et al., 2018). Second, this discrepancy might depend on the genomic regions that were examined. Indeed, we found no clear evidence of newly formed promoter-enhancer loops in response to TNF-α when we re-analyzed our Hi-C data for the *IL1A* and *CITED4* loci, where Jiu et al. examined promoter-enhancer interactions by 3C (data not shown). More recently, circadian gene expression was shown to control rhythmic chromatin interactions between enhancers and promoters within sub-TADs, by *in situ* Hi-C in mouse livers harvested 12 hrs apart (Kim et al., 2018). In addition, promoter-anchored chromatin loops were rapidly reorganized within 4 hrs after inducing the differentiation of 3T3-L1 preadipocytes, as detected using promoter capture Hi-C (Siersbaek et al., 2017). Based on these findings, we believe that chromosomal conformation changes, such as SE-SE loops and enhancer-promoter loops, could be rapidly reorganized at sub-TADs levels even in completely differentiated cells in response to external stimuli, including pro-differentiation and inflammatory signals.

3C-based technologies have been increasingly used to link noncoding GWAS variants associated with inflammatory diseases to distal regulatory elements (Warren et al., 2017). Through genetic fine mapping, Gupta and colleagues identified rs9349379, a common SNP in the third intron of the phosphatase and actin regulatory protein 1 (*PHACTR1*) gene, as the putative causal variant for five vascular diseases, including CAD. In CRISPR-edited stem cell-derived human ECs, the deletion of rs9349379 increased the gene expression of endothelin 1 (*EDN1*), which is located 600 kb upstream of *PHACTR1*. Interestingly, 4C-seq revealed a very low contact probability between rs9349379 and the *EDN1* promoter, but a relatively high contact probability between rs9349379 and the *EDN1* downstream SEs marked with high levels of H3K27ac, suggesting a potential role of rs9349379 in SE function (Gupta et al., 2017). In addition, cardiovascular disease-related SNPs are highly enriched in vascular endothelial growth factor (VEGF)-regulated compartments. One SNP in particular (rs1371799) is located in an enhancer-enhancer loop at the *IL8* locus, where we also confirmed a rapid chromosomal conformation change in response to TNF-α (Kaikkonen et al., 2014). Consistently, our study has revealed that many genetic variants associated with cardiovascular diseases, including CAD, atherosclerosis, hypertension, and stroke, were located in TNF-α-induced SEs that were associated with the recruitment of KDM7A and UTX in human ECs. Thus, the dynamic rewiring of SE-SE chromatin loops in response to inflammation could be an important signal to induce inflammatory gene expression programs leading to cardiovascular diseases.

In summary, our study offers a new perspective on the spatial transcriptional control during inflammatory responses and delineates the multiple mechanisms utilized by KDM7A and UTX in the regulation of the TNF-α-responsive gene expression program in human ECs. Our findings support a model through which KDM7A and UTX mediate the vast majority of the NF-kB signaling pathway, likely via the demethylation of repressive histone marks. The evidence showing that the recruitment of KDM7A and UTX to the inflammatory loci following TNF-α-stimuli is associated with the looping between SEs suggests the possible roles of chromatin interactions in early inflammation. Moreover, we demonstrated that the pharmacological and simultaneous inhibition of KDM7A and UTX markedly reduced leukocyte adhesion *in vivo*. A phase III clinical trial (BETonMACE) is in progress to test the effects of RVX 208, a novel, orally active BET protein inhibitor, in patients with an increased risk of cardiovascular disease (Ghosh et al., 2017). Clearly, histone modification enzymes are ubiquitously expressed in the human body, and therefore cell- or tissue-specific targeting might be required for their translational relevance. Nevertheless, KDM7A and UTX could be novel therapeutic targets for vascular inflammatory diseases, including atherosclerosis.

## ACKNOWLEDGEMENTS

We thank Mika Kobayashi and Shogo Yamamoto (The University of Tokyo) for technical support and bioinformatics analysis, respectively. This work was supported by a Grant-in-Aid for JSPS Postdoctoral Fellows (to Y.H.), a Grant-in-Aid for JSPS Overseas Fellows (to Y.A.), a Grant-in-Aid for Young Scientists (B) 17K15991 (to Y.H.), a Grant-in-Aid for Young Scientists (A) 26710013 (to Y.K.), a Grant-in-Aid for Scientific Research (B) [18H02824 (to M.N.) and 17H03614 (to Y.K.)], a Grant-in-Aid for Scientific Research on Innovative Areas (Research in a proposed research area) 25125707 (to Y.K.), a Grant-in-Aid for Challenging Exploratory Research [26670397 (to Y.K.) and 16K15438 (to Y.K.)], a Fund for the Promotion of Joint International Research (Fostering Joint International Research) 15KK0251 (to Y.K.), AMED CREST [18gm0510018h0106 (to S.T.) and 16gm0510005h0006 (to H.K. and Y.W.)], MEXT KAKENHI 18H05527 (to H.K.), a Research grant by Nanken-Kyoten, TMDU (to Y.H., Y.K., Y.W., and T.F.), a Research grant by Takeda Science Foundation (to Y.K.), a Research grant by Japan Heart Foundation (to Y.K.), a Research grant by MSD Life Science Foundation (to Y.K.), and a Research grant by Kowa Life science Foundation (to Y.K.).

## SUPPLEMENTAL FIGURE LEGENDS

Figure S1. Transcriptional changes of human EC genes in response to TNF-α-stimuli, related to Figure 1

(A-C) Bar plots showing temporal changes of mRNA levels of vascular cell adhesion molecule-1 (VCAM1) (A), intracellular cell adhesion molecule-1 (ICAM1) (B), and E-selectin (SELE) (C) after stimulation of human endothelial cells (ECs) with TNF-α. Data are shown as means ± SD. (D) Time-course western blot analyses for VCAM1, ICAM1, SELE, and β-actin in lysates from ECs treated with TNF-α. (E and F) Gene ontology (GO) analysis of TNF-α-up-regulated genes in class 1 (E) and class 2 (F), as determined in Figure 1A. The *P*-values of each category analyzed by DAVID (https://david.ncifcrf.gov/) are shown in the bar graphs. (G) Heat map representation of the TNF-α-down-regulated genes. Genes were classified based on the TNF-α-treated time points, using the algorithm “HOPACH”. (H) GO analysis of TNF-α-down-regulated genes, as determined in Figure S1G. The *P*-values of each category analyzed from DAVID are shown in the bar graphs. (I and J) Bar plots showing mean mRNA levels of VCAM1 measured 4 hrs after stimulation of ECs with or without TNF-α ± miR-374b-5p (I) and miR-374a-5p (J). Data are shown as means ± SE. \**P* < 0.05 compared to TNF-α (-) mimics (-). † *P* < 0.05 compared to TNF-α (+) mimics (-).

Figure S2. KDM7A and UTX participate in TNF-α-induced NF-κB-p65 signaling pathways, related to Figure 2.

(A-D) Scatter plots showing reproducibility of RNA-seq signals between the two biological replicates in Figure 3A. The R^2^ correlation coefficient is shown in each graph. (E-G) Venn diagrams showing overlap of the genes downregulated by BAY and siKDM7A+siUTX (E), siKDM7A (F), or siUTX (G) treatments of TNF-α-responsive genes (KDM7A: lysine demethylase 7A, UTX: lysine demethylase 6A). The overlapping area shows the number of overlapping genes. (H-K) GO analysis of TNF-α-responsive genes downregulated by BAY (H), siKDM7A+siUTX (I), siKDM7A (J), or siUTX (K) treatments, as determined in Figure S2E-G. The *P*-values of each category analyzed by DAVID are shown in the bar graphs.

Figure S3. KDM7A and UTX control TNF-α-induced expression of adhesion molecules in human ECs, related to Figure 2.

(A-D) Bar plots showing mean mRNA levels of VCAM1 (A), ICAM1 (B), SELE (C), and KDM7A (D) measured 4 hrs after stimulation of ECs with or without TNF-α ± siKDM7A. Data are shown as means ± SE. ^*^*P* < 0.05 compared to TNF-α (-) siKDM7A (-). † *P* < 0.05 compared to TNF-α (+) siKDM7A (-). (E) Western blots for VCAM1, ICAM, SELE, and β-actin in lysates from ECs treated with or without TNF-α (4 hrs) ± siKDM7A. (F) Representative images (top) and bar plot quantification (bottom) showing adhesion of calcein-labeled U937 monocytes to ECs treated with or without TNF-α ± siKDM7A. Scale bar represents 250 μm. Data are shown as means ± SD. \**P* < 0.05 compared to TNF-α (-) siKDM7A (-). † *P* < 0.05 compared to TNF-α (+) siKDM7A (-). (G-J) Bar plots showing mean mRNA levels of VCAM1 (G), ICAM1 (H), SELE (I), and UTX (J) measured 4 hrs after stimulation of ECs with or without TNF-α ± siUTX. Data are shown as means ± SE. ^*^*P* < 0.05 compared to TNF-α (-) siUTX (-). † *P* < 0.05 compared to TNF-α (+) siUTX (-). (K) Western blots for VCAM1, ICAM, SELE, and β-actin in lysates from ECs treated with or without TNF-α (4 hrs) ± siUTX. (L) Representative images (top) and bar plot quantification (bottom) showing adhesion of calcein-labeled U937 monocytes to ECs treated with or without TNF-α ± siUTX. Scale bar represents 250 μm. Data are shown as means ± SD. ^*^*P* < 0.05 compared to TNF-α (-) siUTX (-). † *P* < 0.05 compared to TNF-α (+) siUTX (-).

Figure S4. Characterization of KDM7A-, UTX-, and p65-recruited elements, related to Figure 3.

(A) Western hybridization detection of exogenous KDM7A and UTX in human ECs 1 day after adenoviral infection. (B and C) Scatter plots showing reproducibility of ChIP-seq signals for KDM7A (B) and UTX (C) between the two biological replicates. The R^2^ correlation coefficient is shown in each graph. (D) Pie charts of KDM7A (left) and UTX (right) binding site distributions in ECs with TNF-α (+). (E) Metagene representations of KDM7A (left) and UTX (right) ChIP-seq signals in units of read count per million mapped reads at a meta composite of the genomic regions around TNF-α-down-regulated genes. (F and G) Gene tracks of ChIP-seq signals for KDM7A and UTX, and RNA-seq signals around the *TBX1* (F) and *CALHM2* (G) loci in TNF-α (-) and TNF-α (+) ECs. ChIP-seq and RNA-seq signals are visualized by the Integrated Genome Viewer (http://software.broadinstitute.org/software/igv/). (H) Pie chart of p65 binding site distribution in ECs with TNF-α (+). (I) *De novo* motif analysis of p65 binding sequences in TNF-α-stimulated ECs. The *P*-value indicates significant enrichment of the transcription factor binding motif in p65 binding sites in comparison to a size-matched random background. (J) Boxplots of p65, H3K27ac, KDM7A, and UTX signals at all p65 binding sites in ECs before and after TNF-α stimulation. The Y axis indicates normalized read counts (See STAR Methods). \**P* < 0.05 compared to TNF-α (-).

Figure S5. Characterization of SEs and ChIA-PET interactions, related to Figure 5. (A and B) The average profiles of enrichment of ChIP-seq experiments for bromodomain-containing protein 4 (BRD4; top) and H3K27ac (bottom) under TNF-α treatment (TNF+) and control (TNF-) around broad peaks of the KDM7A (A) or UTX (B) ChIP-seq experiment. (C) Plot of enhancers defined in TNF-α-untreated ECs, ranked by increasing BRD4 signal. Super enhancers (SEs) were defined by ROSE (Loven et al., 2013; Whyte et al., 2013), based on the ChIP-Seq signals for BRD4 (Brown et al., 2014). SEs are indicated by dashed lines and yellow boxes. The numbers denotes the total SEs and their classification based on the binding of KDM7A and UTX. Representative SE-related genes are indicated in the graph. (D and E) ChIA-PET interactions of active RNA pol II around the *SOX18* (D) and *NR2F2* (E) loci integrated with the ChIP-seq profiles of KDM7A and UTX, and the RNA-seq profiles in TNF-α (-) and TNF-α (+) ECs. ChIA-PET interactions were visualized by the WashU Epigenome Browser (http://epigenomegateway.wustl.edu/browser/). Interactions detected by ChIA-PET are depicted with purple lines. Red bars show control-specific SEs.

Figure S6. Characterization of Hi-C data, related to Figure 6.

(A) Fractions of intra- and inter-chromosomal interactions identified by Hi-C that connect regions within valid interactions. Valid interactions or duplicates, and intra- or inter-chromosomal interactions were calculated by HiC-Pro (https://github.com/nservant/HiC-Pro). See also STAR Methods. (B) Violin plots showing reproducibility between all Hi-C replicates. The Y axis indicates Pearson’s correlation coefficient. (C and D) Virtual 4C views and heat maps of Hi-C data around the *PALMD* locus (C) and the *HOXA* cluster (D). Virtual 4C calculated from Hi-C datasets, using bins covering the indicated viewpoints. The Y axis in the virtual 4C view indicates the number of reads that interact with the viewpoint bin. Control reads are raw reads and TNF-α reads were normalized to control reads. Topologically associating domains (TADs) were calculated by TADtool (https://github.com/vaquerizaslab/tadtool) and indicated as blue bars. Red lines indicate SEs.

Figure S7. Single drug effect of the KDM7A or UTX enzymatic inhibitor against monocyte adhesion in mice, related to Figure 7.

Male, 10 week old C57BL/6 mice (n=3/group) were pretreated with vehicle, Daminiozed (50 mg/kg), or GSKJ4 (50 mg/kg) at 16 hrs and 1 hr prior to TNF-α treatment. Inflammation was induced by an intraperitoneal injection of TNF-α (5 μg/mouse) and IVM was performed 4 hrs later. (A) Representative snapshot from IVM analysis of leukocyte adhesive interactions in the femoral arteries. Margins of vessels are indicated with dashed lines. White spots represent fluorescently labeled leukocytes visualized by an intravenous injection of rhodamine 6G. (B) Quantitative analyses of leukocyte adhesive interactions in the femoral arteries. Data are shown as means ± SE. ^*^*P* < 0.05 compared to the vehicle group.

## STAR⋆METHODS

Detailed methods are provided in the online version of this paper and include the following:

- KEY RESOURCES TABLE
- CONTACT FOR REAGENT AND RESOURCE SHARING
- EXPERIMENTAL MODEL AND SUBJECT DETAILS
  - Mice
  - Cell lines
- METHODS DETAILS
  - TNF-α stimulation
  - mRNA and miRNA isolation
  - miRNA-microarray
  - Transfection procedure
  - Real-time PCR for mRNA quantification
  - Western blot
  - Adhesion assay
  - RIP followed by microarray
  - Luciferase assay
  - Cloning of KDM7A and UTX
  - Adenovirus infection
  - Intravital microscopy
  - ChIP
  - ChIP-qPCR
  - ChIP-seq library preparation
  - RNA-seq library preparation
  - ChIA-PET library preparation
  - *In situ* Hi-C library preparation
  - Bioinfomatics
- QUANTIFICATION AND STATISTICAL ANALYSIS
- DATA AND SOFTWARE AVAILABILITY

## KEY RESOURCES TABLE

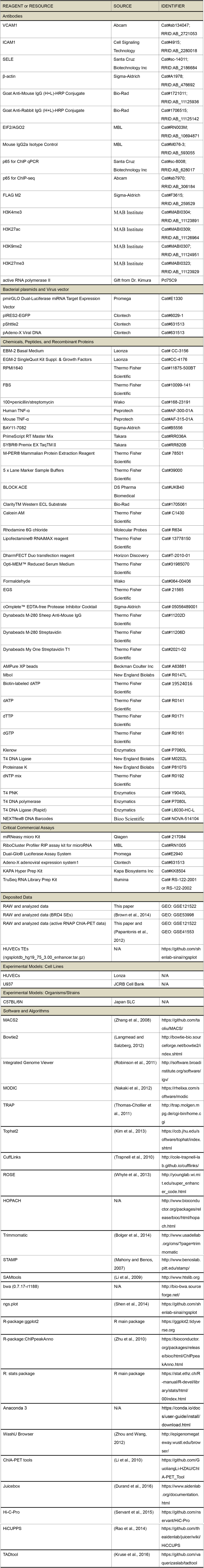

## CONTACT FOR REAGENT AND RESOURCE SHARING

Further information and requests for resources and reagents should be directed to, and will be fulfilled by the Lead Contact, Yasuharu Kanki.

## EXPERIMENTAL MODEL AND SUBJECT DETAILS

Sources of cell lines and mouse models used in the study are reported in the Key Resources Table. All mouse experiments were approved by The University of Tokyo Animal Care and Use Committee.

### Mice

Male C57BL/6N mice aged 10 weeks were purchased from Japan SLC (Shizuoka, Japan). The animals were housed in individual cages in a temperature- and light-controlled environment and had *ad libitum* access to chow and water.

### Cell lines

Human Umbilical Vein Endothelial Cells (HUVECs) were used as human ECs in this work. HUVECs were purchased from Lonza Japan Ltd. (Tokyo, Japan) and cultured and passaged every 2 or 3 days in EBM-2 complete medium composed of EBM-2 Basal Medium (Lonza), supplemented with EGM-2 SingleQuot Kit Suppl. & Growth Factors (Lonza) and 3% fetal bovine serum (FBS) (Thermo Fisher Scientific, Waltham, MA). We sub-cultured HUVECs when they reached 80 to 90% confluence, to avoid excessive floaters. HUVECs were used within passages 4 to 7 for all experiments in this work. The human monocyte cell line U937 was purchased from the JCRB Cell Bank (Osaka, Japan) and grown and passaged every 2 or 3 days in RPMI1640 (Thermo Fisher Scientific), supplemented with 1% penicillin/streptomycin (Wako, Osaka, Japan) and 10% FBS. All cells were cultured at 37 °C and in a 5% CO_2_ atmosphere in a humidified incubator.

### Methods Details

#### TNFα stimulation

HUVEC plates (80-90% confluent) were washed with PBS once and then serum starved in EBM-2 Basal Medium containing 0.5% FBS for 16 hrs. HUVECs were stimulated with or without 10 ng/ml TNF-α (Peprotech, Rocky Hill, NJ) for the indicated periods thereafter.

#### mRNA and miRNA isolation

Total RNA, including miRNA, was isolated from HUVECs using a miRNeasy micro Kit (Qiagen, Hilden, Germany) with the DNase digestion step, according to the manufacturer’s instructions.

#### miRNA-microarray

The miRNA-microarray (Human miRNA Microarray, Release 19.0, 8×60K; Agilent Technologies, Santa Clara, CA) was carried out by MBL (Aichi, Japan).

#### Transfection procedure

Synthetic miRNA mimics for miR-374b-5p, miR-374a-5p, and miR-3679-5p, negative control miRNA mimics, as well as inhibitors for miR3679-5p and negative control inhibitors were obtained from Thermo Fisher Scientific. siRNAs for lysine demethylase 7A (KDM7A) and 6A (UTX), and negative control siRNA were purchased from Thermo Fisher Scientific or Sigma-Aldrich (St. Louis, MO). Sequences of miRNA mimics and siRNAs are listed in Table S5. MiRNA mimics, miRNA inhibitors, or siRNAs were transfected with Lipofectamine^®^ RNAiMAX reagent (Thermo Fisher Scientific) according to the manufacturer’s protocol. Growth medium was replaced 5 hrs after transfection.

#### Real-time PCR for mRNA quantification

The isolated RNA (500 ng) was reverse-transcribed to cDNA with PrimeScript RT Master Mix (Takara, Shiga, Japan). PCR was performed with a CFX96 unit (Bio-Rad, Hercules, CA) with SYBR^®^ Premix EX Taq^TM^II (Takara). Relative expression levels were calculated using β-actin mRNA as a reference. The primers for quantification are listed in Table S5.

#### Western blot

Whole cells were lysed in M-PER^®^ Mammalian Protein Extraction Reagent (Thermo Fisher Scientific), and then the lysate was centrifuged at 10,000 x g for 10 min to pellet the cell debris. The supernatant was mixed with 5 x Lane Marker Sample Buffer (Thermo Fisher Scientific), and then incubated for 5 min at 95°C. Proteins were separated by SDS-PAGE and then transferred to nitrocellulose membranes (Bio-Rad). After blocking with 4% BLOCK ACE blocking agent (DS Pharma Biomedical, Osaka, Japan), the membrane was incubated with the following antibodies: monoclonal rabbit anti-vascular cell adhesion molecule-1 (VCAM1) (1:2,500 dilution) (Abcam, Cambridge, MA), polyclonal rabbit anti-intracellular cell adhesion molecule-1 (ICAM1) (1:2,500 dilution) (Cell Signaling Technology, Danvers, MA), polyclonal rabbit-E-selectin (SELE) (1:1,000 dilution) (Santa Cruz Biotechnology Inc., Santa Cruz, CA), and mouse monoclonal anti-β-actin (1:5,000) (Sigma-Aldrich) at 4°C overnight. The blots were then incubated with a peroxidase-conjugated anti-rabbit or mouse IgG antibody (1:10,000 dilution) (Bio-Rad) at 37°C for 40 min. The antibody-antigen reaction was detected with the Clarity^TM^ Western ECL Substrate (Bio-Rad).

#### Ahesion assay

Monocyte adhesion to HUVECs was assayed as previously described (Inoue et al., 2006; Tozawa et al., 2011). In brief, HUVECs were grown to confluence in a 24-well cell culture plate. U937 cells were labeled with the fluorescent dye Calcein AM (Thermo Fisher Scientific) in EBM2 containing 0.5% FBS. HUVECs transfected with miRNA mimics or siRNAs were incubated with or without TNF-α (10 ng/ml) for 4 hrs, and then U937 cells were added to the plate in a final concentration of 3 x 10^5^ cells per well. After 1 hr, the cells were washed twice with Hanks’ balanced salt solution containing calcium and magnesium (Thermo Fisher Scientific). Adhesion of U937 cells was visualized with a fluorescent microscope. Nine microscopic fields under each condition were photographed at random, using a Leica DMI6000 B camera with an Adaptive Focus Control system (Leica Microsystems K.K. Tokyo, Japan), and then the U937 cells were automatically counted with the ImageJ 1.48v program (https://imagej.nih.gov/ij/download.html).

#### RIP followed by microarray

RNA immunoprecipitation (RIP) was performed using a RiboCluster Profiler RIP assay kit for microRNA (#RN1005; MBL), according to the manufacturer’s protocol. Briefly, HUVECs transfected with miR-3679-5p or negative control miRNA mimics were stimulated with TNF-α and then immunoprecipitated with magnetic Dynabeads M-280 (Thermo Fisher Scientific) coupled with an anti-EIF2C2/AGO2 mouse monoclonal antibody (MBL). A mouse IgG2a Isotype control antibody (MBL) was used as a negative control to confirm the proper functioning of the RIP assay (data not shown). Large RNAs and small RNAs were separately extracted from the antibody immobilized beads-RNP complex, and then the AGO2-bound large RNAs were subjected to the microarray analysis.

Preparation of cRNA and hybridization of probe arrays were performed according to the manufacturer’s instructions (Affymetrix, Santa Clara, CA, USA). Affymetrix Genechip Human Genome U133 plus 2.0 arrays containing over 54,000 probe sets were used. The expression value for each mRNA was obtained with the Affymetrix Microarray Suite 5 method (MAS5). To analyze the expression data at the genetic level, the probes with the highest expression value for each gene were selected, and subsequently the probes exhibiting a value of more than 1,000.0 in at least one condition were screened. Finally, probes with an average fold change (miR-3679-5p/control) greater than 2.0 or less than 0.5 were extracted.

#### Luciferase assay

The 3′-UTRs of the human KDM7A and UTX mRNAs were amplified by PCR from the cDNA of HUVECs and cloned at the *Sac*1-*Xba*1 sites for KDM7A and the *Sac*1-*Xho*1 sites for UTX into the pmirGLO Dual-Luciferase miRNA Target Expression Vector (Promega, Madison, WI). The PCR primers and oligonucleotide sequences for the constructs are listed in Table S5. All of the constructs were further confirmed by sequencing. For the luciferase activity analysis, each construct was co-transfected with the miRNA mimics in a 12-well plate, using the DharmFECT Duo transfection reagent (Horizon Discovery, Cambridge, UK) for 24 hrs, and the luciferase assays were performed with the Dual-Glo Luciferase assay system (Promega), according to the manufacturer’s protocol. The transfection efficiency for each well was normalized by the Renilla luciferase activity.

#### Cloning of KDM7A and UTX

Human full length KDM7A was produced by ligating the fragment of 1-382 bp obtained by synthesis (Thermo Fishers Scientific) and the 382-2816 bp fragment cloned by PCR using cDNA from HUVECs. UTX was cloned by PCR using cDNA from the pluripotent human testicular embryonal carcinoma cell line, NTERA-2 cl.D1 (American Type Culture Collection, Manassas, VA). The PCR primers and oligonucleotide sequences for the constructs are listed in Table S5. The cDNA was subcloned into pIRES2-EGFP (Clontech, Shiga, Japan) and then transferred into the pShuttle2 and pAdeno-X Viral DNA (Clontech) by using the Adeno-X adenoviral expression system1 (Clontech). All cloned constructs were confirmed by restriction enzyme digestions and DNA sequencing.

#### Adenovirus infection

We preliminarily performed a setup experiment to test different multiplicities of infection (MOI), and an MOI of 100 was considered to be optimum (data not shown). Therefore, we applied an MOI of 100 for further analysis in this study. One day before the transduction, 1.0 ×10^6^ HUVECs were seeded into 15 cm dishes in 15 ml of EBM-2 complete medium. The HUVECs exhibited 80 to 90 % confluence at the time of transduction. Adenovirus particles were mixed with 2 ml of Opti-MEM^™^ Reduced Serum Medium (Thermo Fisher Scientific) and incubated for 30 min at room temperature. HUVECS were washed with PBS once and then incubated with 2 ml of the adenovirus containing Opti-MEM^™^ Reduced Serum Medium for 1 hr at 37 °C in a 5% CO_2_ atmosphere in a humidified incubator. Afterwards, 15 ml of EBM-2 complete medium was added to the culture dish.

#### Intravital microscopy

Male, 10 week old C57BL/6N mice (n=3-4/group) were pretreated with vehicle (5% DMSO), Daminozide (50 mg/kg), GSKJ4 (50 mg/kg), or mixed inhibitor (Daminiozed+GSKJ4, 50 mg/kg each) at 16 hrs and 1 hr prior to TNF-α treatment. Inflammation was induced by an intraperitoneal injection of mouse recombinant TNF-α (PeproTech) at a dose of 5 μg/mouse. Leukocyte adhesive interactions in the femoral arteries were assessed by intravital microscopy (IVM), as previously described (Ito et al., 2016; Osaka et al., 2007). In brief, mice were injected with rhodamine 6G chloride (Molecular Probes, Eugene, OR) via the right femoral vein to label leukocytes *in vivo*. The left femoral artery was visualized with a microscope (model BX51WI; Olympus, Tokyo, Japan) equipped with a water immersion objective. Labeled leukocytes were clearly visualized in the anterior half of the vessels facing the objective, and platelets less than 5 μm in diameter were excluded. All images were recorded using a personal computer with an image analysis program (MetaMorph; Molecular Devices, Sunnyvale, CA). The numbers of adherent and rolling leukocytes (*i.e*., those that did not move for &#x2267; 3 s during the 1 min recording period and those that passed a reference line perpendicular to the vessel axis, respectively) were counted along a region of interest, a 100×100 μm segment of the vessel, and expressed as the number of interacting cells per 10^4^ μm^2^ of the vessel surface.

#### ChIP

Chromatin immunoprecipitation (ChIP) was performed as previously described (Kanki et al., 2011; Kanki et al., 2017). In brief, 10 million HUVECs were cross-linked with 1% formaldehyde (Wako) for 10 min. For the p65 and FLAG ChIP analyses, cells were fixed with 2 mM EGS (ethylene glycol bis [succinimidyl succinate]) (Thermo Fisher Scientific) for 1 hr prior to formaldehyde fixation. After neutralization by using 0.2 M glycine, the cells were collected, re-suspended in 2 ml SDS lysis buffer, composed of 10 mM Tris-HCl, pH 8.0 (Thermo Fisher Scientific), 150 mM NaCl (Thermo Fisher Scientific), 1% SDS (Sigma-Aldrich), 1 mM EDTA, pH 8.0 (Thermo Fisher Scientific), and cOmplete^™^ EDTA-free Protease Inhibitor Cocktail (Sigma-Aldrich), and fragmented by a Picoruptor (40 cycles for histones and p65, and 20 cycles for FLAG, 30 sec on/30 sec off; Diagenode, Liege Science Park, Belgium). The sonicated solution was diluted with ChIP dilution buffer (20 mM Tris-HCl, pH 8.0, 150 mM NaCl, 1 mM EDTA, 1% Triton X-100 (Sigma-Aldrich)) up to 10.3 ml, and 10 ml were used for immunoprecipitation (10 ml) and the remaining 300 μl were saved as non-immunoprecipitated chromatin (INPUT). The specific antibody was bound to magnetic Dynabeads M-280 and applied to the diluted, sonicated solution for immunoprecipitation. Antibodies against H3K4me3, H3K27ac, H3K9me2, H3K27me3 (MAB Institute, Inc. Hokkaido, Japan), p65 (Abcam for ChIP-Seq and Santa Cruz Biotechnology Inc. for ChIP-PCR), and FLAG (Sigma-Aldrich) were used. The prepared DNA was quantified using a NanoDrop 2000 spectrophotometer (Thermo Fisher Scientific), and more than 10 ng of DNA were processed for qPCR or sequencing.

#### ChIP-qPCR

The primers for quantification are listed in Table S5. PCR was performed with a CFX96 PCR and SYBR® Premix EX Taq^TM^II. Fold enrichment was determined as the percentage of the input.

#### ChIP-seq library preparation

For the ChIP-seq analysis using the FLAG antibody, the precipitated DNA from the original DNA size of ~2 kb was further sheared to ~200 bp by an Acoustic Solubilizer (Covaris, Woburn, MA). For ChIP-seq using an antibody against histones, DNA sonicated to an average size of 0.5 kb was used for the ChIP-seq library preparation. ChIP samples were processed for library preparation with a KAPA Hyper Prep Kit (Kapa Biosystems Inc., Wilmington, MA), according to the manufacturer’s instructions. Deep sequencing was performed on a HiSeq 2500 sequencer (Illumina Inc., San Diego, CA) as single-end 50 base reads.

#### RNA-seq library preparation

Total RNA from the cells was isolated as described above. The RNA integrity score was calculated with the RNA 6000 Nano reagent (Agilent Technologies) in a 2100 Bioanalyzer (Agilent Technologies). All samples used for the preparation of the RNA-seq libraries had a RIN (RNA Integrity Value) score above 9. RNA-Seq libraries were prepared with a TruSeq RNA Library Prep Kit (Illumina). The libraries were sequenced on a HiSeq 2500 system (Illumina) as paired-end 150 base reads.

#### ChIA-PET library preparation

Chromatin interaction analysis with paired-end tag sequencing (ChIA-PET) was performed as previously described (Li et al., 2010; Papantonis et al., 2012). Briefly, cells were crosslinked using 10 mM EGS in 50% glacial acetic acid (45 min) and then in 1% paraformaldehyde (20 min), quenched (5 min) in 2.5 M glycine, harvested, and sonicated (Digital Sonifier 250; Branson, Danbury, CT). The ChIP was performed using magnetic beads (Dynabeads M-280) with the Pd75C9 antibody directed against phospho-Ser2/-Ser5 within the heptad repeat at the C-terminus of the largest catalytic subunit of RNA polymerase II (a gift from H. Kimura). The chromatin captured on the magnetic beads was trimmed to create blunt ends, phosphate groups were added to the 5′ ends, the ends were ligated to each other via biotinylated half-linkers, and the complexes were eluted. After the crosslinking was reversed, the DNA was purified and digested with *Mme*I (binding site encoded by the linker). After immobilization on M-280 Streptavidin Dynabeads (Thermo Fisher Scientific), the adaptors were ligated, and the efficiency of library production was evaluated by PCR. Finally, the di-tags were prepared, the library was sequenced on a GAII analyzer (Illumina), and the resulting paired-end tags (PETs) were analyzed.

#### *In situ* Hi-C library preparation

*In situ* Hi-C was performed in two replicates for HUVECs treated with or without TNF-α for 60 min. Cells were cross-linked for 10 min at room temperature with 1% formaldehyde and quenched for 5 min at room temperature with 0.2 M glycine. The cross-linked cells were centrifuged at 2,500 x g for 5 min at 4°C. To isolate nuclei, cross-linked cells were resuspended in 200 μl of lysis buffer (10 mM Tris-HCl (pH 8.0), 10 mM NaCl, 0.2% NP40, Protease Inhibitor Cocktail (Sigma-Aldrich)) and incubated on ice for 15 min. The suspension was then centrifuged at 2,500 x g for 5 min, and the pellet was washed by resuspension in 300 μl of lysis buffer and centrifugation at 2,500 x g for 5 min at 4°C. The pellet was resuspended in 50 μl of 0.5% SDS and incubated for 10 min at 62°C. After heating, 170 μl of 1.47% Triton X-100 was added to the suspension and incubated for 15 min at 37°C. To digest the chromatin, 25 μl of 10X NEBuffer2 and 100U *Mbo*I (New England Biolabs, Beverly, MA) were added to the suspension and incubated at 37°C overnight with rotation. Enzymes were inactivated by heating for 20 min at 62°C. Fragmented ends were biotin-labeled by adding 50 μl of a mixture containing 0.3 mM biotin-14-dATP (Thermo Fisher Scientific), 0.3 mM dATP (Thermo Fisher Scientific), 0.3 mM dTTP (Thermo Fisher), 0.3 mM dGTP (Thermo Fisher Scientific), and 0.8 U/ul Klenow (Enzymatics, Beverly, MA) and incubating the solution for 1 hr at room temperature with rotation. The biotin-labeled fragmented ends were subsequently ligated by adding 900 μl of a mixture containing 120 μl of 10X T4 DNA ligase buffer (New England Biolabs), 100 μl of 10% Trition X-100, 12 μl of 10 mg/ml BSA (New England Biolabs), 5 μl of 400 U/μl of T4 DNA Ligase (New England Biolabs), and 663 μl H_2_O, and incubating the solution for 4 hrs at room temperature with rotation. The nuclei were pelleted for 5 min at room temperature at 2,500 x g. After removing the supernatant, the nuclei were resuspended in 550 μl of 10 mM Tris-HCl (pH 8.0). The nuclei were reverse-crosslinked by adding 50 μl of Proteinase K (New England Biolabs) and 57 μl of 10% SDS, and incubating the mixture for 30 min at 55°C. Subsequently, the nuclei were mixed with 67 μl of 5 M NaCl and incubated at 68°C overnight. After cooling the samples for 10 min at room temperature, the DNA was captured with 0.8X AMPureXP beads (Beckman Coulter Inc., Brea, CA) and eluted in 100 μl of 10 mM Tris-HCl (pH 8.0). DNA shearing was performed using a focused-ultrasonicator (Covaris, E220; Power 140 W, Duty factor 10%, Cycle per bust 200, Time 67 sec, temperature 7°C) and the final sample volume after shearing was brought to 200 μl with 10 mM Tris-HCl (pH 8.0). To design the DNA selection for 200-600 bp size, a 120 μl aliquot of AMPureXP beads was added to 200 μl of sample (beads ratio: 0.6) and incubated for 5 min at room temperature with inversion. To remove the DNA fragments >600 bp, the clear supernatant was collected on a magnet and the beads were discarded. As a second round of size selection, an 80 μl aliquot of AMPureXP beads was added to the supernatant and incubated for 5 min at room temperature with inversion. The supernatant was removed on a magnet, and the beads were washed twice with 700 μl of 70% ethanol and dried for 5 min. DNA fragments (200-600 bp) were eluted from the beads with 300 μl of 10 mM Tris-HCl (pH 8.0). For the biotin pull down, a 100 μl aliquot of 10 mg/ml Dynabeads My One T1 Streptavidin beads (Thermo Fisher Scientific) was washed with 1X Tween Wash Buffer (5 mM Tris-HCl (pH 7.5), 0.5 mM EDTA, 1 M NaCl, 0.05% Tween). The beads were resuspended in 300 μl of 2X Binding Buffer (10 mM Tris-HCl (pH 7.5), 1 mM EDTA, 2M NaCl), transferred to sample tubes, and incubated for 15 min at room temperature. The beads were subsequently washed twice with 1X Tween Wash Buffer for 2 min at 55°C with mixing and washed once with 1X NEB T4 DNA ligase buffer (Enzymatics). To repair the fragmented ends and remove the biotin from unligated ends, the beads were resuspended in 85 μl of 1X NEB T4 DNA ligase buffer (Enzymatics), 5 μl of 10 mM dNTP Mix (Thermo Fisher Scientific), 5 μl of 10 U/μl NEB T4 PNK (New England Biolabs), 4 μl of 3 U/μl NEB T4 DNA Polymerase (New England Biolabs), and 1 μl of 5 U/μl Klenow (Enzymatics), and incubated for 30 min at room temperate. After the supernatant was discarded on a magnet, the beads were washed twice with 1X Tween Wash Buffer for 2 min at 55°C with mixing and resuspended in 100 μl of 1X NEB Buffer2. For dA-tailing, the beads were resuspended in 90 μl of 1X NEB Buffer2, 5 μl of 10 mM dATP (Thermo Fisher Scientific), and 5 μl of 5 U/μl Klenow (3ʹ→5ʹ exo-) (Enzymatics), and incubated for 30 min at 37°C. The beads were subsequently washed twice with 1X Tween Wash Buffer as before and once with 1X NEB Quick Ligation Reaction Buffer (New England Biolabs). To ligate the adapters, the beads were suspended in 50 μl of 1X NEB DNA Quick Ligation Reaction Buffer, 1 μl of NEXTflex® DNA Barcodes (Bioo Scientific, Austion, TX), and 2 μl of NEB DNA Quick Ligase (New England Biolabs), and incubated for 15 min at room temperature. The beads were washed twice with 1X Tween Wash Buffer as before and once with 10 mM Tris-HCl (pH 8.0), and resuspended in 20 μl of 10 mM Tris-HCl (pH 8.0). After estimating the concentration and the cycle number for the final PCR by a qPCR assay, the final PCR was directly performed with the T1 beads, using a KAPA Hyper Prep Kit (KAPA Biosystems Inc.). The DNA was cleaned with 1X AMPure beads, eluted in 30 μl of 10 mM Tris-HCl (pH 8.0), and analyzed by paired-end sequencing.

### Bioinformatics

#### RNA-seq data analysis

Sequence reads (36-bp single read) were aligned to the human reference genome (GRCh37/hg19) with Tophat (version: 2.1.1) (Kim et al., 2013) with the following parameters: the mismatch to the reference genome should be supported by at least 2 reads, and the lengths of insertions and deletions should be supported by at least 3 bp. After assigning the mapped reads onto the gene positions deposited in the geocode database (https://www.gencodegenes.org/releases/19.html), the FPKMs (fragments per kilobase of exon per million reads) of all of the deposited genes were calculated by CuffLinks (Trapnell et al., 2010) with the default parameters. RNA-seq signals were visualized with the Integrated Genome Viewer (Version 2.4.8) (http://software.broadinstitute.org/software/igv/). The RNA-seq signal of each locus was normalized by the following basis:

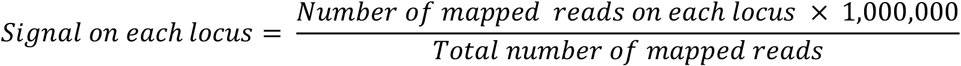

#### Reproducibility between RNA-seq experiments

The reproducibility of the genome-wide RNA-seq signals in the biological replicates was examined under all conditions. The FPKM values were used as the RNA-seq signals. Subsequently, the correlation coefficients between two biological replicates were calculated, based on the FPKMs of each reference gene.

#### Gene Ontology Analysis

The gene annotation enrichment analysis was performed for the Gene Ontology (biological process and cellular component), using the Functional Annotation tool at DAVID Bioinformatics Resources 6.7 (http://david.abcc.ncifcrf.gov/).

#### PCA analysis

The first three eigenvectors of the six samples (*i.e*., control, TNF-α, TNF-α with BAY 11-7082, siKDM7A, siUTX, and siKDM7A+siUTX) were used to analyze the expression patterns of TNA-α-responsive genes. In total, 404 TNF-α-upregulated genes (more than 2-fold after TNF-α treatment in comparison to the control) and 136 TNF-α-downregulated genes (less than 0.5-fold after TNF-α treatment in comparison to the control) were used as the TNF-α-responsive genes in the principal component analysis (PCA). The variances of each eigenvector are 74.9% for PC1, 9.6% for PC2, and 6.4% for PC3, respectively.

#### ChIP-Seq data analysis

Sequence reads (36-bp single read) from ChIP for histones, KDM7A, UTX, and p65 were mapped to the human reference genome hg19, by using the Illumina alignment program, ELAND. Only sequence reads that mapped uniquely to the reference genome were used for further analysis. The putative binding sites (peaks) of KDM7A and UTX were analyzed by SICER (version: 1.1) with the default parameter settings (Window size: 200 bp, Gap size: 400 bp, E-value: 100). The binding signals of the ChIP-seq for KDM7A and UTX were constructed by piling up the read counts onto the whole genome, in which the size of each sequence read was commonly set to be 2,000 bp. The putative binding sites (peaks) of p65 were analyzed by MACS (version 1.4.2) (Zhang et al., 2008) with the default parameter settings. The binding signals of the ChIP-seq for p65 were obtained by using the calibration of the read counts by MACS. The ChIP-seq signals were visualized by the Integrated Genome Viewer (Version 2.4.8) (http://software.broadinstitute.org/software/igv/).

#### Reproducibility between ChIP-seq experiments

The reproducibility of the genome-wide ChIP-seq signals for KDM7A and UTX was examined by comparing two biological replicates. First, the ChIP-seq peaks of the two biological replicates were merged. Subsequently, the correlation coefficients between the two biological replicates were calculated, based on the count of the mapped sequence reads of the merged peaks.

#### Distribution of ChIP-seq signals

The ChIP-seq signals of H3K27ac, p65, KDM7A, and UTX in human ECs with or without TNF-α treatment were compared in the TNF-specific binding sites of KDM7A or UTX. In each binding site, the 10 kb regions from the centers of the KDM7A and UTX binding sites were screened, and the ChIP-seq signals were normalized by the total read counts.

#### Genomic Regions Enrichment of Annotations Tool (GREAT)

GREAT version 3.0.0 (http://great.stanford.edu/public/html/index.php) was performed on the KDM7A- and UTX-binding regions obtained from the ChIP-seq results, in order to identify their functional annotations (McLean et al., 2010).

#### *De novo* motif analysis

The *de novo* binding motifs were identified by the MODIC motif identification program (window size: 12 bp, background sequence: random genomes, enrichment ratio: >2.0 fold change compared to random genomes) (Nakaki et al., 2012). The ten sequence motifs enriched in each ChIP-seq assay were outputted by the program and integrated into three motifs, based on the sequence patterns. Specifically, the sequence pattern of each motif was converted into a position weight matrix, and the motifs with correlation coefficients > 0.8 between two matrices were integrated to the significant one. The three *de novo* motifs were assigned to the known TF-binding motifs deposited in TRANFAC (Matys et al., 2003) (http://genexplain.com/transfac/) by using the STAMP web tool (Mahony and Benos, 2007) (http://www.benoslab.pitt.edu/stamp/).

#### Boxplot

The p65 binding sites before and after TNF-α stimulation were merged as the set of the total p65 binding sites. The set includes three patterns (*i.e*., control-specific, common, and TNF-α-specific) of p65 binding. The normalized read counts for each site of the set were calculated for the ChIP-seq analyses of p65, H3K27ac, KDM7A, and UTX before and after TNF-α stimulation. To determine whether a significant difference exists between binding before and after TNF-α stimulation, a Students t-test was used to compare the binding distributions of both treatments after normalization. The normalized read counts were calculated as follows:

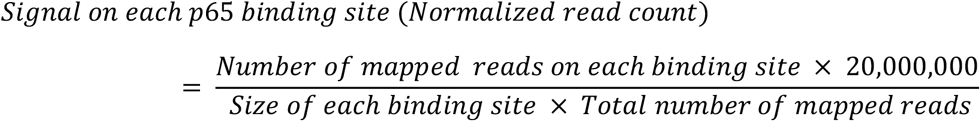

#### Prediction of super enhancers in TNF-treated conditions

Super enhancers (SEs) were predicted using the ROSE software package, as described in (Loven et al., 2013; Whyte et al., 2013) and available at (younglab.wi.mit.edu/super_enhancer_code.html). First, the BRD4 ChIP-seq data were downloaded from GSE54000 (https://www.ncbi.nlm.nih.gov/geo/query/acc.cgi?acc=GSE54000) and the binding sites were analyzed as described in the section “ChIP-seq data analysis”. Subsequently, super enhancers were identified by ROSE. The putative super enhancers were ranked, based on the BRD4-binding intensities and the special enrichment of BRD4-binding sites. Comparing the BRD4-binding sites in HUVECs with or without TNF-α treatment, the super enhancers were classified into 3 groups: TNF-α-specific, control-specific, and common. The sites with more than 1-bp overlapping between TNF-α 0 min and 60 min are regard as “common” super enhancers.

#### Scatter plot

To visualize the effects of TNF-α stimulation on the gene recruitments by KDM7A (Figure 3A) and UTX (Figure 3B), the log2 fold changes of mRNA-seq expression levels (horizontal axis) and KDM7A- or UTX-ChIP-seq binding levels (vertical axis) between TNF-α-stimulated and non-TNF-α-stimulated samples were plotted. The genes were categorized by the log2 fold changes as the TNF-α-up-regulated genes (red; FC > 1,5), TNF-α-down-regulated (blue; FC < 1.5), and unchanged (grey; other).

#### NGS plot

To compare the effects of TNF-α stimulation on the recruitment of KDM7A and UTX, the average ChIP-seq expression levels of KDM7A (Figure 3C) and UTX (Figure 3D) around the transcription start sites (TSS) of target genes were plotted using an NGS plot (Shen et al., 2014), where the target genes were selected as the TNF-α-up-regulated and down-regulated genes (up: FC > 4, down: FC < 2).

The effects of TNF-α stimulation on the recruitments between the SEs and typical enhancers (TEs) were compared for KDM7A (Figure 5D) and UTX (Figure 5E) via a comparison of the ChIP-seq expression levels after the TNF-α stimulations around the SE and TE regions, using an NGS plot. The SEs in Figure 5D and 5E included all of the control-specific, common, and TNF-α-specific SEs as defined above. TEs were downloaded from the pre-compiled HUVECs ngsplotdb (ngsplotdb_hg19_75_3.00_enhancer.tar.gz) (https://github.com/shenlab-sinai/ngsplot).

#### Analyses of repressive histone modification scores

The enrichment of ChIP-seq reads (H3K9me2, H3K27me3, and their input DNAs) in gene bodies with TNF-α treatment (TNF+) and control (TNF-) was quantified in Reads Per Kilobase Million (RPKM). The log2 fold change of ChIP enrichment, compared with its input DNA, was then computed for each of the KDM7A and UTX target genes and shown by box plots (Figures 4A and 4B). The KDM7A and UTX target genes were selected as follows: (1) the log2 fold change of mRNA-seq expression levels with TNF-α treatment (TNF+) and control (TNF-) was larger or equal to 2; (2) the log2 fold change of ChIP signal between a ChIP experiment (KDM7A or UTX) and its input DNA was larger or equal to 0.5.

#### ChIA-PET data processing

ChIA-PET libraries yielded ~35×10^6^ 20-bp paired-end reads each, from which 10.8×10^6^ and 8.8×10^6^ were successfully aligned to the genome (hg19) for the 0-, 30-, and 60-min samples, respectively. For stringency, two or more reads with the same sequences, or mapping within 2 bp of the left and right ends of another read, were classified as one PET.

#### Analysis of ChIA-PET interaction changes

The range of each binding site was set to 10 kb. The number of ChIA interactions that started on each binding site was normalized, using the total ChIA count for each condition. For each binding site, the rate of the interaction count was plotted as Bean plots or Pirate plots.

#### Hi-C data processing

The paired read data obtained for the Hi-C libraries were processed with the HiC-Pro (Servant et al., 2015) pipeline, using the human genome (hg19) and the fragment information for the restriction enzyme “*Mbo*I”. Finally, 196,814,231 and 204,936,615 valid pairs of reads remained for the control and TNF-α-treated samples, respectively. The Hi-C raw data were converted to a Juicebox compatible format (.hic format), using the Juicer-Tools software.

#### Contact matrices

The raw Hi-C data of chromosomes 1 and 4 were exported using Juicer-Tools with 1 Mb, 50 kb, and 3 kb bin sizes. Interaction scores were visualized by a color gradient, as shown in Fig. 6A and B.

#### Virtual 4C

Virtual 4C data show the interaction counts starting at viewpoints (within 5 kb), which are extracted from Hi-C data. Those viewpoint-containing interactions were counted by 10 kb bins to either end position. The count-data is then shown as a line graph. The virtual 4C data of TNF-α were normalized by those of the control (*i.e*., no TNF-α treatment).

#### Hi-C loop calling (HiCCUPS)

The significant Hi-C loops were calculated using the Juicer Tools software with the following option “hiccups –m 512 –r 5,000, 10,000, 25,000-f 0.1, 0.1, 0.1-p 4, 2, 1-i 7, 5, 3-d 20,000, 20,000, 50,000”.

#### TAD boundary calling

The raw Hi-C matrices with 10 kb resolution were used for TAD calling. TADs were obtained by the TADtool algorithm (Kruse et al., 2016), with 100,000 bp as the window size and 10 as the cut off.

#### SEs overlap with GWAS data

Single nucleotide polymorphisms (SNPs) satisfying genome-wide significance for coronary artery disease (CAD), atherosclerosis, hypertension, or stroke were downloaded from the NHGRI-EBI GWAS Catalog of published genome-wide association studies (GWAS) (Welter et al., 2014). SNPs having strong linkage disequilibrium (LD) (r^2^ >= 0.8) to the reported GWAS SNPs (CAD, atherosclerosis, hypertension, and stroke) in the European reference population of the 1000 Genomes Project (Genomes Project et al., 2015) were extracted by querying the HaploReg v3 database (http://www.broadinstitute.org/mammals/haploreg/haploreg.php). We extracted SNPs that are located on the 356 EC-specific SEs defined in Figure 5A from among the SNPs detected in the HaploReg v3 database, by overlapping their physical positions (hg19 genome build).

## QUANTIFICATION AND STATISTICAL ANALYSIS

Statistical differences were analyzed by the Tukey–Kramer test for multiple comparisons. Differences between two groups were compared by the Student’s *t* test. In all tests, differences with *P* values of <0.05 were considered statistically significant.

## DATA AND SOFTWARE AVAILABILITY

The array and sequence data can be accessed through the Gene Expression Omnibus (GEO) under the NCBI accession number GSE121522.

## SUPPLEMENTAL ITEMS

Table S1. Summary of analysis for RNA-seq, miRNA microarray, and RIP followed by microarray, related to Figure 1.

Table S2. Summary of siRNA experiments, related to Figure 2.

Table S3. Summary of GREAT analysis, related to Figure 3.

Table S4. Summary of GWAS results, related to Figure 5.

Table S5. List of miRNAs, siRNAs, and primers used for this study.

Movie S1, IVM experiment in TNF-α-treated mice, related to Figure 7.

Movie S2, IVM experiment in TNF-α-treated mice with Daminozide and GSK-J4, related to Figure 7.

